# Structure-guided allosteric modulation of the delta opioid receptor

**DOI:** 10.1101/2025.10.16.682975

**Authors:** Jesse I. Mobbs, M. Deborah Nguyen, Owindeep Deo, Damian Bartuzi, Hariprasad Venugopal, Sadia Alvi, Vi Pham, Nick Barnes, Arthur Christopoulos, Daniel P. Poole, Simona E. Carbone, Manuela Jörg, Ben Capuano, Jens Carlsson, Arisbel B. Gondin, Peter J. Scammells, Celine Valant, David M. Thal

## Abstract

Opioid analgesics remain essential for pain management but are associated with significant adverse effects, including respiratory depression, tolerance, and dependence. The δ-opioid receptor (δOR) represents a promising therapeutic target for developing safer opioid analgesics with reduced adverse effects compared to conventional μ-opioid receptor-targeting drugs. Positive allosteric modulators (PAMs) offer advantages over direct agonists by enhancing endogenous opioid signaling while preserving natural spatiotemporal activation patterns, potentially avoiding tolerance and dependence issues. Here, we present high-resolution cryo-EM structures of δOR complexed with the peptide agonist DADLE and the PAM MIPS3614, revealing a novel lipid-facing allosteric binding site formed by transmembrane helices 2, 3, and 4. MIPS3614 stabilizes the active receptor conformation through a critical hydrogen bond with residue N131^3.35^ in the conserved sodium binding site, a key regulatory region controlling GPCR activation. Comprehensive mutagenesis, molecular dynamics simulations, and structure-activity relationships validate this proposed mechanism. Structure-guided optimization yielded MIPS3983 with enhanced binding affinity and retained cooperativity. Our findings establish the first molecular framework for δOR allosteric modulation and provide a structural foundation for the rational design of safer opioid therapeutics.

## Introduction

Opioid analgesics are essential for pain management but are associated with significant adverse effects, including respiratory depression, gastrointestinal dysfunction, tolerance, and dependence^1^. With prescription-based drug dependence already beyond crisis point, the prevalence of these safety issues has contributed to the current opioid crisis, emphasizing the need for therapeutic approaches with improved safety profiles.

Opioid receptors comprise a family of four G protein-coupled receptors (GPCRs): mu (μ), delta (δ), kappa (κ), and nociceptin/orphanin FQ peptide (NOP) receptors, which predominantly couple to inhibitory G_i_ proteins^2^. While μ-opioid receptor (µOR) activation produces both the beneficial analgesia and detrimental side effects of conventional opioids^3,4^, the δ-opioid receptor (δOR) has emerged as a promising alternative therapeutic target^5^. δOR agonists demonstrate efficacy in preclinical models of chronic pain, migraines, and mood disorders with reduced respiratory depression and dependence liability compared to µOR agonists^6–10^. Despite this promise, first-generation δOR agonists targeting the orthosteric binding site encountered development challenges due to on-target side effects, including seizures and tolerance development^11–13^. While next-generation δOR agonists have successfully mitigated seizure risk, they still face significant hurdles, including the development of tolerance, insufficient clinical efficacy, and off-target effects^10,14^.

δOR agonists also represent promising therapeutic targets for gastrointestinal (GI) disorders. The enteric nervous system expresses functional δOR, and activation of the δOR-enkephalin axis provides important inhibitory control over intestinal motility and secretion^15^. This is evidenced clinically by racecadotril, an enkephalinase inhibitor that enhances endogenous opioid signaling to treat secretory diarrhea^16^. The ability of positive allosteric modulators (PAMs) to selectively enhance endogenous opioid signaling when and where it occurs naturally makes them particularly attractive for GI applications, potentially providing therapeutic benefit for conditions like irritable bowel syndrome with diarrhea (IBS-D) while preserving physiological control mechanisms and avoiding the adverse effects associated with sustained orthosteric receptor activation.

Despite these potential advantages across multiple therapeutic areas, a fundamental challenge for δOR agonists stems from their mechanism of action, which involves direct orthosteric site activation that drives receptor desensitisation and tolerance^12^. Alternative approaches to safer opioid therapeutics have explored ligands with distinct pharmacological profiles^17^. These include receptor-targeted strategies such as partial agonists that exhibit reduced intrinsic efficacy^18^, biased agonists that selectively activate beneficial signaling pathways^19–21^, allosteric modulators that target distinct allosteric sites to enhance endogenous ligand activity without directly activating the receptor^23,24^, and bitopic ligands that bind to both orthosteric and allosteric sites, conferring both efficacy and selectivity to a single chemical entity^25,26^. Additionally, indirect approaches include enkephalinase inhibitors that prevent degradation of endogenous opioid peptides and potentially avoid tolerance development^22^.

PAMs targeting δOR represent a promising alternative therapeutic strategy^27–29^. In response to pain, the body releases endogenous peptides such as enkephalins and endorphins, which activate opioid receptors to provide localized, natural pain relief. However, during severe or chronic pain, endogenous opioids are often insufficient, requiring administration of analgesics. Rather than directly activating the receptor, PAMs bind to distinct allosteric sites and selectively enhance the activity of endogenous opioid peptides, thereby preserving natural spatiotemporal signaling patterns while potentially circumventing some limitations inherent to orthosteric agonists^30^. The first discovered δOR PAM, BMS-986187, potentiates enkephalin-mediated responses in a model system^23^. However, its poor physicochemical properties have limited its utility to peripheral *in vivo* studies, where it has shown potential in treating GI motility disorders^15^. In structural biology studies, BMS-986187’s poor solubility likely prevented the determination of its allosteric site in X-ray crystallography studies^31^. We report a similar null finding when using BMS-986187 to determine a DADLE-bound δOR-G_i_ complex by cryo-EM. Fortunately, a medicinal chemistry campaign exploring the structure-activity relationships (SAR) of the xanthenedione scaffold of BMS-986187 has yielded MIPS3614, which exhibits substantially reduced cLogP values, resulting in improved compound solubility while maintaining similar pharmacological properties to those of BMS-986187^32^.

In this study, we present a high-resolution cryo-EM structure of a δOR-DADLE-G_i_ complex co-bound to the PAM MIPS3614, revealing a novel lipid-facing allosteric binding site formed by residues on transmembrane helices 2, 3, and 4. Our structural analysis reveals that the carbonyl oxygen of MIPS3614 forms a hydrogen bond with residue N131^3.35^, which constitutes part of the conserved allosteric sodium ion binding site^33,34^. Molecular dynamics simulations demonstrate that MIPS3614 stabilizes an outward, lipid-facing rotamer conformation of N131^3.35^ specifically associated with the receptor’s active state. We validate this mechanism through comprehensive mutagenesis and functional studies, demonstrating that removal of the carbonyl oxygen completely abolishes PAM activity and confirming the critical importance of the N131^3.35^ interaction. We demonstrate that MIPS3614 effectively modulates δOR signaling in both recombinant systems and native tissues, providing proof-of-concept for therapeutic potential in GI disorders. Through structure-guided optimisation targeting additional binding site residues, we developed enhanced δOR PAMs, including MIPS3983, which exhibits improved binding affinity while maintaining cooperativity in recombinant systems. Collectively, our findings identify a previously uncharacterised allosteric binding site and elucidate the molecular mechanism of δOR PAMs, establishing a framework for the rational design of this potential new class of therapeutics.

## Results

### Structure of δOR-DADLE

Several structures of the delta opioid receptor (δOR) were determined in complex with small molecule agonists and antagonists, as well as exogenous selective peptides^26,31,33,35–38^, while limited molecular characterization of endogenous peptide δOR agonists exists. We first determined the structure of δOR in complex with the agonist DADLE ([D-Ala^2^, D-Leu^5^]-Enkephalin, Tyr-D-Ala-Gly-Phe-D-Leu). DADLE is a synthetic analogue of the endogenous peptide hormone Leu-Enkephalin (Tyr-Gly-Gly-Phe-Leu), both of which have similar pharmacological and selectivity profiles for the δOR^39^.

To obtain structures of the δOR G protein complex, we used a chimeric miniature G protein (mG_si_), similar to one previously used^40^, where the mG_si_ was fused to the carboxyl terminus of the receptor. This construct was co-expressed with Gβ_1_ and Gγ_2_ subunits, and complex formation was initiated with DADLE. The δOR-mG_si_-DADLE complex was further stabilized by the addition of Nb35^41^ and purified in detergent by anti-FLAG affinity and size exclusion chromatography. Similar to previous studies^37^, we observed anti-parallel oligomers of the δOR complex but were able to separate the monomeric complex for structure determination via size exclusion chromatography (**Supplementary Fig. 1**). The purified δOR-mG_si_-DADLE complex was vitrified on gold grids and imaged using single-particle cryo-transmission electron microscopy (cryo-EM) on a Titan Krios microscope (**Supplementary Table 1**). We determined the cryo-EM structure of δOR-mG_si_-DADLE to a global resolution of 2.0 Å (**Fig. 1a-c, Supplementary Fig. 2 and Table 1**), which was sufficient to model the majority of the backbone and side chains for the receptor, G protein, Nb35, and DADLE (**Supplementary Fig. 3a**).

**Fig. 1.**
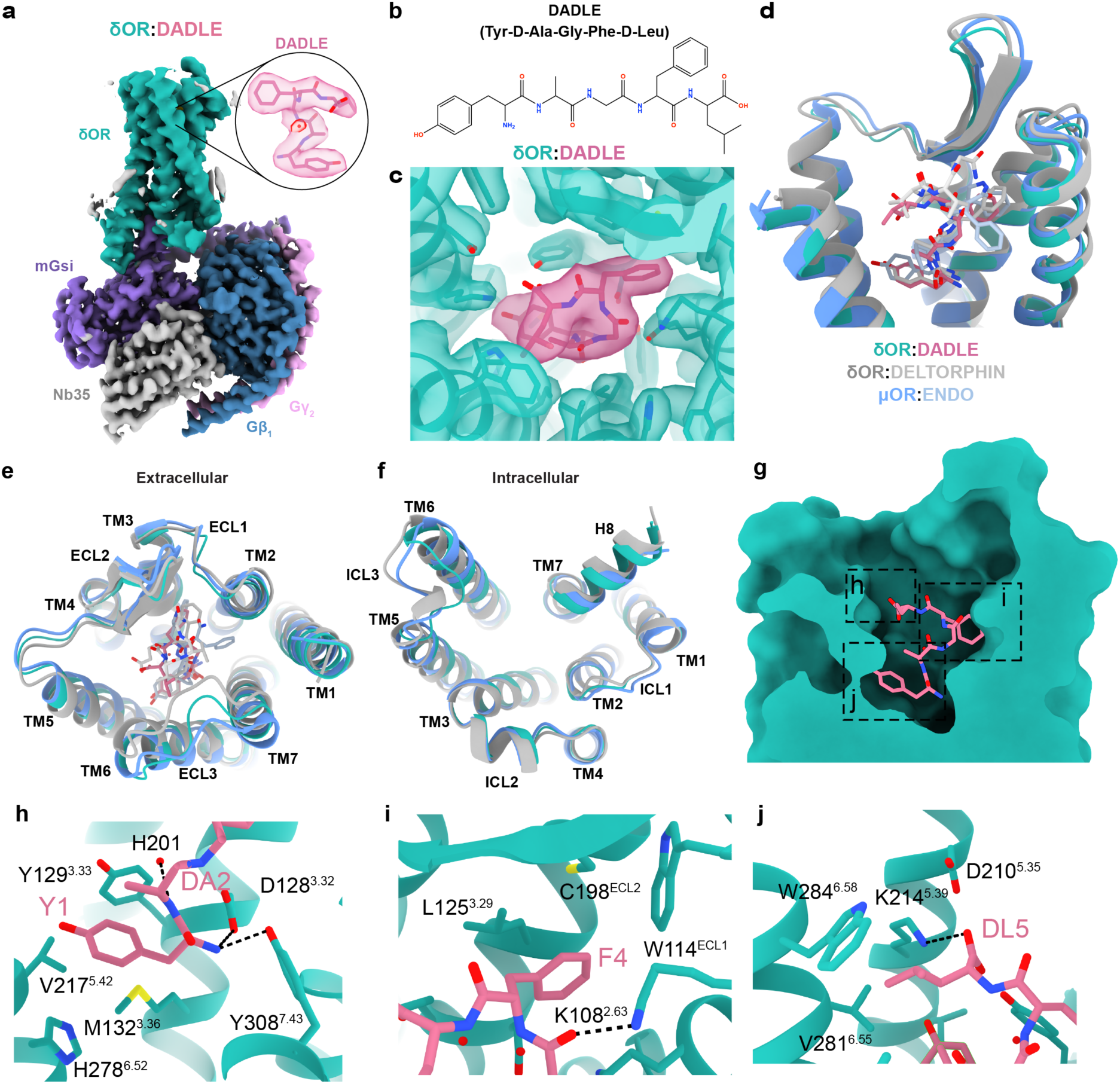
Overall structure of δOR-DADLE complex and comparison with related opioid receptor structures. **(a)** Cryo-EM structure of the δOR-DADLE-mG_si_ complex at 2.0 Å resolution. The δOR is shown in teal, DADLE peptide in pink, mG_si_ in purple, Gβ_1_ in dark blue, Gγ_2_ in pink, and Nb35 in grey. The inset shows a detailed view of DADLE bound in the orthosteric binding site with cryo-EM density contoured at 0.27. (**b**) Chemical structure of DADLE (Tyr-D-Ala-Gly-Phe-D-Leu), a synthetic analogue of the endogenous peptide Leu-Enkephalin. (**c**) Cryo-EM density map contoured at 0.27 showing the quality of density for DADLE and surrounding receptor residues, with the boxed region highlighting the peptide binding site. (**d**) Overlay comparison of δOR-DADLE (teal) with δOR-Deltorphin (grey, PDB: 8F7S) and µOR-Endorphin (blue, PDB:8F7R) showing structural similarities. (**e)** Extracellular view comparison of δOR-DADLE with δOR-Deltorphin and μOR-Endomorphin (blue, PDB:8F7R structures), highlighting differences in ECL3. (**f**) Intracellular view showing G protein coupling interfaces and conformational differences, particularly in intracellular loop 3 (ICL3) between the different opioid receptor-peptide complexes. (**g**) Cross-section view of DADLE binding in the orthosteric pocket showing the peptide’s extended conformation with the boxed region highlighting regions of the peptide binding site. (**h-j**). Molecular interactions of (h) the N-terminal tyrosine (Y1) and D-alanine (DA2), (i) phenylalanine residue (F4), and (j) C-terminal D-leucine (DL5) of DADLE with the receptor.

Compared to previous structures of δOR-Deltorphin (PDB: 8F7S)^37^, δOR-DADLE had a root mean square deviation (RMSD) of 1.04 Å for α-carbons across the receptor and 1.51 Å across the entire complex (excluding the dimer in 8F7S). δOR-DADLE was remarkably similar to μOR-Endomorphin (PDB: 8F7R)^37^ with an RMSD of 0.76 Å for α-carbons across the receptor (1.34 Å across the complex). The extracellular pocket of δOR-DADLE was wider compared to δOR-Deltorphin, due to a more open conformation of ECL2 and ECL3. While the intracellular side showed similar receptor and G protein conformations, except for a change in conformation of ICL3 (**Fig. 1d–f**). G protein binding interfaces were similar, with only a modest rotation of the G protein. Some of the observed differences could be attributed to different G protein stabilisation strategies used, or to the neighbouring dimeric interfaces observed in previous complexes (**Supplementary Fig. 4a–g**). Similar to the previous cryo-EM structures, δOR-DADLE adopts a fully active conformation, with all activation motifs (toggle switch, PIF, NPxxY, and DRY) displaying active conformations (**Supplementary Fig. 4h–k**).

The δOR-DADLE structure revealed that DADLE binds deep within the orthosteric binding site and occupies a similar location to that observed in previous structures of δOR-Deltorphin (Tyr-D-Ala-Phe-Glu-Val-Val-Gly) and μOR-Endomorphin (Tyr-Pro-Trp-Phe) (**Fig. 1g-j**)^37^. The side chain of the N-terminal tyrosine (Y1), which is conserved across opioid peptides, sits in a hydrophobic pocket near TMs 3/5/6, surrounded by residues Y129^3.33^, M132^3.36^, V217^5.42^ and H278^6.52^, while the backbone nitrogen forms hydrogen bonds to D128^3.32^ and Y308^7.43^ (**Fig. 1h**). The side chain of F4 of DADLE extends towards a pocket composed of residues of TMs 2/3 and ECL1 and ECL2. Specifically, F4 forms hydrophobic interactions with L125^3.59^, W114^ECL1^ and C198^ECL2^; this region of the peptide is further stabilized by a hydrogen bond from the backbone oxygen of G3 to K108^2.63^ (**Fig. 1i**). The terminal D-leucine (DL5) extends towards TMs 3/4/5/6, and the side chain forms hydrophobic interactions with W284^6.58^, and the carboxyl terminus of DADLE forms a salt bridge to K214^5.39^ (**Fig. 1j**).

### Molecular Pharmacology of δOR PAMs

In a recent SAR study^32^, we discovered a novel analogue of BMS-986187, MIPS3614 (**4**), that had reduced cLogP values and considerably improved solubility (**Fig. 2a,g**). To understand the allosteric effects of BMS-986187 and MIPS3614 in different signaling pathways^18^, increasing concentrations of the allosteric ligand were co-added to increasing concentrations of DADLE in human embryonic kidney (HEK) 293 cells expressing human δOR (**Fig. 2a-l**). The operational model of allosterism^42^ was fit to the data to estimate allosteric parameters of functional cooperativity (*αβ*), PAM affinity (p*K*_B_), and efficacy of the PAMs (*τ*_B_). In cAMP inhibition assays (**Fig. 2b,h**), both PAMs exhibited robust agonism due to high signal amplification, which made it difficult to fit an allosteric model to the data. In contrast, in less amplified assays, evidenced by significantly lower potencies of DADLE in these assays compared to cAMP, such as β-arrestin 2 (**Fig. 2c,i**), mG_si_ (**Fig. 2d,j**), and Nb33 (**Fig. 2e,k**) recruitment, both PAMs displayed less agonism, and the data could be quantified with an allosteric model (**Fig. 2f,l**). Both PAMs exhibited similar levels of functional cooperativity (*αβ* = 5-7) and low levels of PAM agonism (*τ*_B_ =0.1–1) across all recruitment assays. However, the functional binding affinity of MIPS3614 was approximately 10-fold lower (p*K*_B_ = 4.5-4.8) versus BMS-986187 (p*K*_B_ = 5.7-5.9), suggesting that the improvements made to PAM solubility were at the cost of reduced binding affinity.

**Fig. 2.**
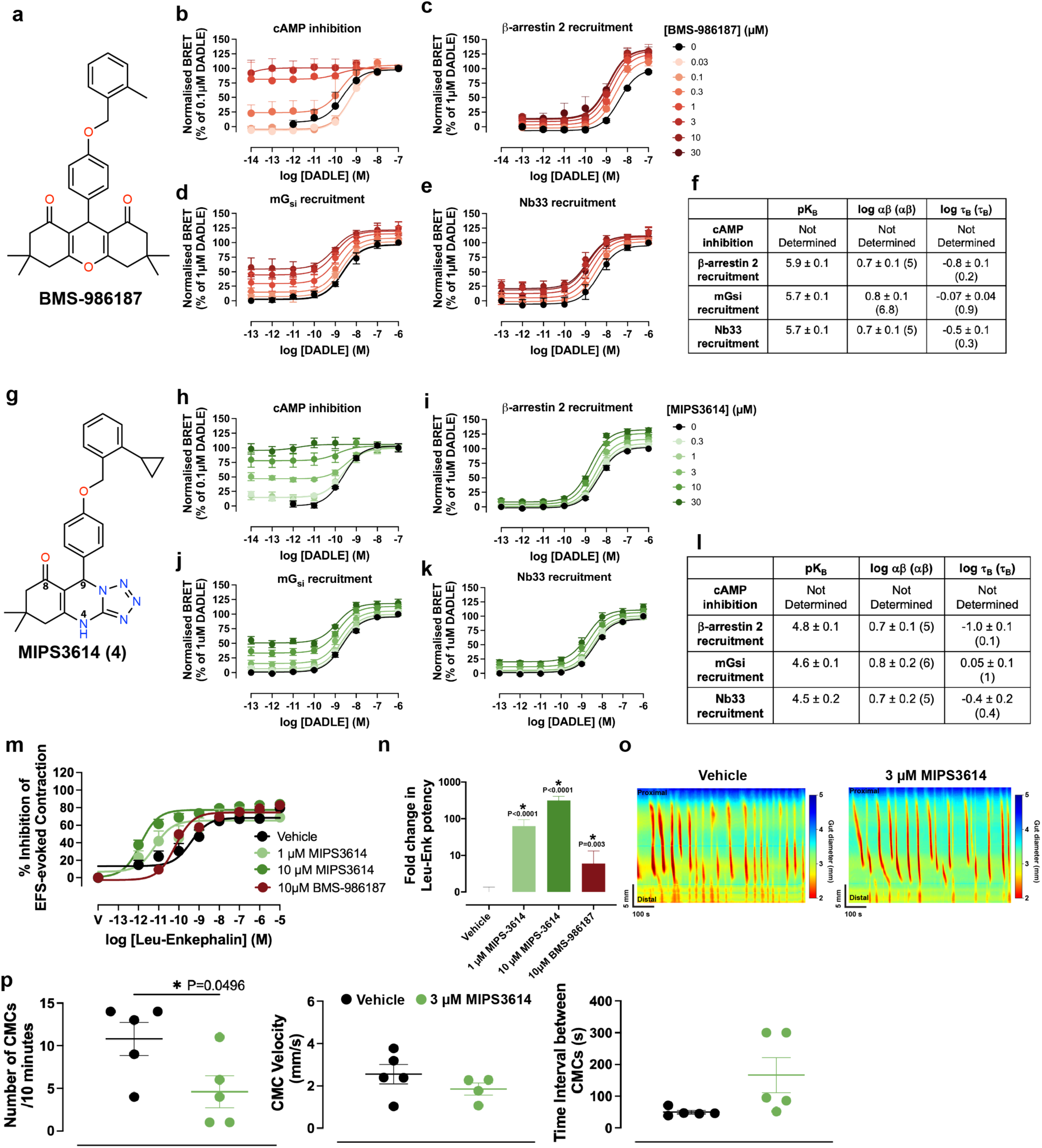
Molecular pharmacology and functional characterisation of δOR positive allosteric modulators BMS-986187 and MIPS3614. **(a)** Chemical structure of BMS-986187, a first-generation δOR PAM with limited solubility. (**b-f**) Pharmacological characterisation of BMS-986187 across multiple signaling pathways. Concentration-response curves showing DADLE-mediated responses in the presence of increasing concentrations of BMS-986187 (0-30 μM) in: (b) cAMP inhibition assay (n=2-3), (c) β-arrestin 2 recruitment assay (n=4-5), (d) mG_si_ recruitment assay (n=3-5), and (e) Nb33 recruitment assay (n=3-4). (f) Summary table of allosteric parameters derived from fitting the operational model of allosterism to the data in (b-e) to estimate functional cooperativity (*αβ*), PAM affinity (p*K*_B_), and PAM efficacy (*τ*_B_) values. (**g**) Chemical structure of MIPS3614, a second-generation δOR PAM with improved solubility compared to BMS-986187 through reduced cLogP values. Key atoms are numbered. (**h-l**) Equivalent pharmacological characterisation of MIPS3614 using the same assay panel as BMS-986187 (h) cAMP inhibition assay (n=5), (i) β-arrestin 2 recruitment assay (n=6), (j) mG_si_ recruitment assay (n=5), and (k) Nb33 recruitment assay (n=5). (l) A summary table shows that MIPS3614 exhibits similar functional cooperativity and low intrinsic efficacy compared to BMS-986187, but with approximately 10-fold lower binding affinity. (**m**) Validation of MIPS3614 allosteric effects in native tissue. Concentration-response curves for Leu-Enkephalin-mediated inhibition of electrically evoked contractions of the mouse colon in the absence (vehicle) or presence of 1 μM MIPS3614, 10 μM MIPS3614, or 10 μM BMS-986187. Data for 10 μM BMS-986187 were reproduced from Deo *et al*., 2025 for comparison^32^. (**n**) Quantification of Leu-Enkephalin potency changes from (m), demonstrating that MIPS3614 has a 10-fold greater allosteric effect compared to BMS-986187 in native tissue. Statistical significance was determined with one-way ANOVA (*p < 0.05) with a Dunnett’s post-test using Vehicle for comparison. (**o**) Representative traces from whole-organ colonic motility assays showing colonic motor complexes (CMCs) under vehicle control (left) and 3 μM MIPS3614 treatment (right) with conditions under elevated intraluminal pressure. (**p**) Quantitative analysis of colonic motility parameters showing the effects of 3 μM MIPS3614. MIPS3614 significantly reduced CMC frequency without affecting velocity under conditions of elevated intraluminal pressure. Statistical significance was determined with unpaired two-tailed t tests (*p < 0.05). (b-p) Data are presented as mean ± SEM.

To assess if MIPS3614 had similar activity on the enteric nervous system (ENS) as BMS-986187^15,32^, we examined its allosteric modulation on Leu-Enk-mediated inhibition of electrically evoked contractions of the mouse colon (**Fig. 2m**). MIPS3614 at 1 μM and 10 μM significantly increased Leu-Enk potency by 100-fold and 300-fold, respectively (**Fig. 2n**). Compared to BMS-986187, this corresponds to a 10-fold greater allosteric effect. We then examined 3 µM MIPS3614 in whole-organ colonic motility assays, measuring the frequency and velocity of colonic motor complexes (CMCs) (**Fig. 2o**). Under basal conditions, MIPS3614 had no effect on CMC parameters (data not shown). In contrast, when intraluminal pressure was elevated to increase CMC frequency, MIPS3614 significantly reduced this response and increased the time interval between contractions (**Fig. 2p**) without affecting CMC velocity. Collectively, these data suggest that, like BMS-986187, MIPS3614 can potentiate the effects of endogenous opioids released in response to increased intraluminal pressure.

### Allosteric site of the delta opioid receptor

To determine structures of δOR-bound to allosteric ligands, the δOR-mG_si_-DADLE complex protein was incubated with BMS-986187 or MIPS3614 prior to vitrification and cryo-EM data collection. We resolved 3D reconstructions of δOR-DADLE-BMS-986187 and δOR-DADLE-MIPS3614, both to a global resolution of 1.9 Å (**Fig. 3a-c, Supplementary Figs. 5 and 6**). However, compared to the δOR-mG_si-_DADLE complex structure, we did not observe sufficient cryo-EM density to confidently model BMS-986187 (**Fig. 3b**), a finding consistent with prior attempts to determine the structure using X-ray crystallography^31^. Therefore, subsequent structural analysis was limited to δOR-DADLE and δOR-DADLE-MIPS3614 complexes. The cryo-EM maps for both structures were of high quality (**Supplementary Fig. 3b**). The δOR-DADLE, BMS-986187, and MIPS3614 structures were remarkably similar, with RMSDs of 0.10 – 0.45 Å for α-carbons of the entire complex (**Fig. 3d-f**).

**Fig. 3.**
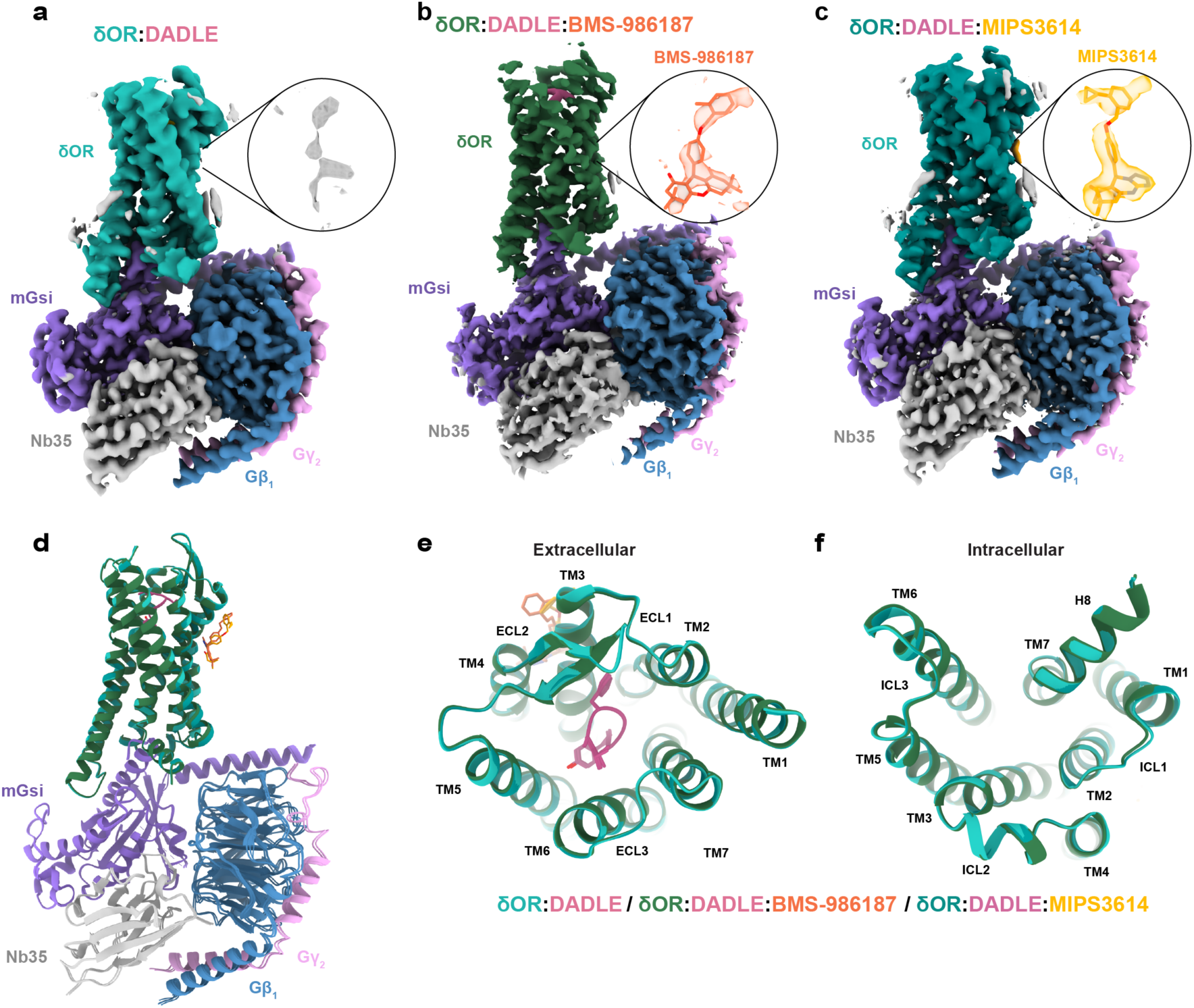
Structural comparison of δOR-DADLE complexes with and without positive allosteric modulators. (**a-c**) Consensus cryo-EM maps for the structures of (a) the δOR-DADLE complex, (b) the δOR-DADLE-BMS-986187 complex, and (c) the δOR-DADLE-MIPS3614 complex contoured at 0.27, 0.45, and 0.30, respectively. δOR is coloured in shades of green, DADLE peptide in pink, mG_si_ in purple, Gβ_1_ in dark blue, Gγ_2_ in pink, and Nb35 in gray. The inset shows cryo-EM density surrounding the allosteric site for MIPS3614 with cryo-EM density from the receptor-focused maps coloured at (a) 0.30, (b) 0.34, and (c) 0.35. (**d-f**) Comparison of the δOR structures from (a-c) with views from the (d) side, (e) extracellular surface, and (f) intracellular surface.

Our structure of the δOR-DADLE-MIPS3614 revealed that the allosteric binding site is located on the extrahelical surface of TMs 2/3/4 (**Fig. 4a-c**). The cryo-EM density surrounding the *ortho*-cyclopropylbenzyl substituent was less defined, suggesting a degree of conformational heterogeneity in this region. Nonetheless, MIPS3614 could be unambiguously modelled (**Fig. 4b**). Notably, all compounds were synthesised as racemic mixtures; however, only the *R*-enantiomer clearly fit the cryo-EM density (**Supplementary Fig. 7a**). Subsequently, chiral HPLC separation of the racemic mixture (MIPS3614) into **4a** and **4b** revealed that only one enantiomer had pharmacological activity, which we have defined as the *R*-enantiomer (**4a**) to match the structural data (**Supplementary Fig. 7b**).

**Fig. 4.**
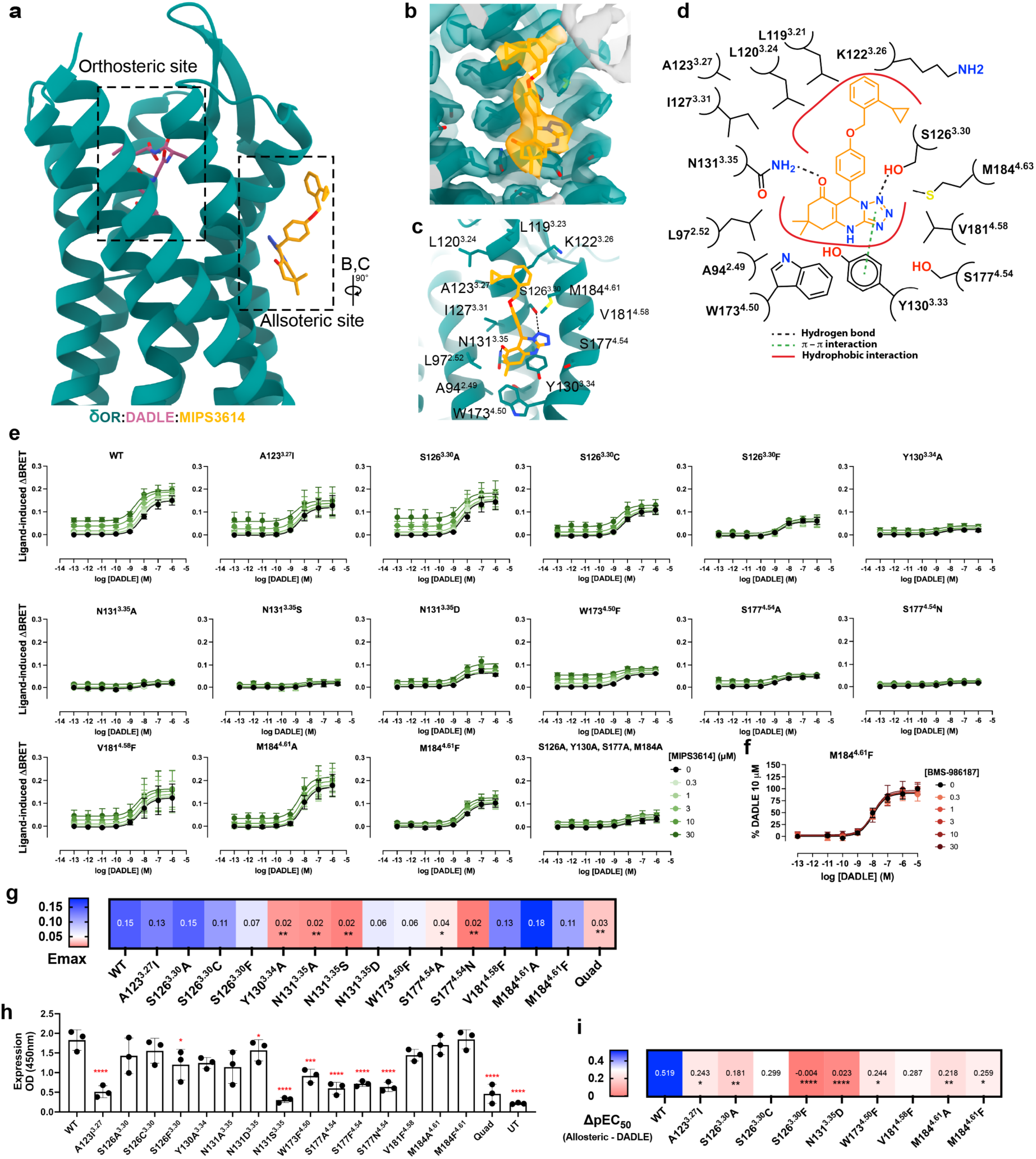
Structural characterization and molecular validation of the MIPS3614 allosteric binding site. **(a)** Overview of the δOR-DADLE-MIPS3614 complex showing the spatial relationship between the orthosteric binding site (DADLE in pink, dashed box) and the allosteric binding site for MIPS3614 in orange (dashed box). The allosteric site is located on the lipid-facing extrahelical surface of transmembrane helices 2, 3, and 4. (**b**) View of the MIPS3614 allosteric binding site with the receptor-focused cryo-EM map contoured at 0.35. (**c)** Detailed view of the MIPS3614 binding site showing key interacting residues from TM2, TM3, and TM4. (**d**) Molecular interaction diagram of MIPS3614 with δOR residues. Hydrogen bonds are denoted by black dashed lines, π-stacking interactions with green dashed lines, and hydrophobic interactions with solid red lines. (**e**) Mutational analysis of the MIPS3614 binding site using the Nb33 recruitment assay. Data show ligand-induced BRET responses. N=3 replicates were performed for all mutant δOR cell lines and n=7 for WT. (**f**) Effect of BMS-986187 at the M184^4.61^F mutant. Data was normalised to 10 µM DADLE response. (**g**) Heat map summarising the effects of mutations on DADLE maximum response (E_max_) from (e). Blue indicates increased E_max_ compared to wild-type, while red indicates reduced E_max_. E_max_ mean values are overlaid on the heatmap. (**h**) Surface receptor expression quantified from ELISA. UT = Untransfected cells. Quad = S126^3.30^A/ Y130^3.34^A/S177^4.54^A/M184^4.61^A. Data are from n=3 experiments for all cell lines. (**i**) Heat map showing the allosteric modulation capacity (Δp*EC*_50_) of MIPS3614 across functional mutants. Values represent the change in DADLE potency in the presence of 30 μM MIPS3614. Red indicates loss of allosteric modulation, while blue indicates retained modulation. Δp*EC*_50_ mean values are overlaid on the heatmap. (e-i) Data are presented as mean ± SEM. Statistical significance was determined by one-way ANOVA with a Dunnett’s post-test using WT for comparison (*p < 0.05, **p < 0.01, ***p < 0.001, ****p < 0.0001). Mean ± SEM and p-values for (g-h) are reported in Supplementary Table 2.

MIPS3614 adopts an extended binding pose with numerous molecular interactions (**Fig. 4c,d**). The tricyclic heterocyclic core is positioned at the centre of the transmembrane region, extending upward toward the extracellular surface. The core sits in a hydrophobic pocket formed by residues L97^2.52^, A94^2.49^, W173^4.50^ and V181^4.68^. Within this pocket, the tricyclic heterocyclic core forms a π-stacking interaction with Y130^3.34^, which is further stabilized by two hydrogen bonds: one from the carbonyl oxygen in the 8-position to N131^3.35^ and another from N1 to S126^3.30^. The *ortho*-cyclopropylbenzyl group primarily interacts through a patch of hydrophobic residues, including L119^3.23^, L120^3.24^, A123^3.27^, I127^3.31^, and M184^4.61^.

To validate the observed interactions, we performed a series of single-point mutations as well as a combined four-residue mutation and tested them in the Nb33 BRET assay. The maximal response (E_max_) of DADLE was significantly reduced at multiple mutants, including Y130^3.34^A, N131^3.35^A, N131^3.35^S, S177^4.54^A, S177^4.54^N, and the quadruple mutant S126^3.30^A/Y130^3.34^A/S177^4.54^A/M184^4.61^A, compared to WT (**Fig. 4e**). The reduction in E_max_ correlated with reduced surface expression levels (**Fig. 4g-h**). Mutation of W173^4.50^, which is conserved across nearly all class A GPCRs, to phenylalanine also resulted in reduced surface expression. In contrast, mutations at positions S126^3.30^, V181^4.58^, and M184^4.61^ were well tolerated without affecting cell surface expression or the DADLE E_max_ response (**Fig. 4g-h**). For mutants that did not significantly reduce DADLE E_max_, we calculated the change in DADLE EC_50_ in the absence or presence of 30 μM of MIPS3614 (**Fig. 4i**). In WT δOR, 30 μM of MIPS3614 caused a 3-fold leftward shift in the DADLE potency, but this potentiation was reduced or absent in most mutants, indicating that residues within this pocket, particularly S126^3.30^, N131^3.35^ and M184^4.61^, are critical for allosteric interactions with MIPS3614.

Mutation of the hydrogen-bonded S126^3.30^ to alanine was well tolerated in terms of surface expression, with DADLE displaying a similar E_max_ to WT, but resulted in a significant reduction of the leftward potency shift of DADLE in the presence of 30μM MIPS3614 (1.5-fold). Mutation of S126^3.30^ to the bulky phenylalanine slightly reduced expression and DADLE E_max_ but completely abolished MIPS3614 modulation. We also mutated M184^4.61^ to an alanine and phenylalanine, both of which reduced the modulation of MIPS3614. Although we were unsuccessful in observing clear cryo-EM density for BMS-986187, the M184^4.61^F mutation had an even greater effect on its binding and modulation (**Fig. 4f**), indicating BMS-986187 binds the same allosteric site as MIPS3614.

Sequence alignment of δOR residues that interact with MIPS3614 with the corresponding μOR and κOR residues reveals the binding site is largely conserved across the opioid family (**Supplementary Fig. 8**). The majority of the site is fully conserved, including residues that form major interactions, such as S126^3.30^ and N131^3.35^, which form hydrogen bonds, and Y130^3.34^, which forms pi-stacking interactions. The differences between the opioid subtypes are primarily subtle changes to hydrophobic residues lining the pocket, such as L119^3.23^I/V (μOR/κOR residue, respectively), A123^3.27^I/I, V181^4.58^L/I and M184^4.61^M/I (**Supplementary Fig. 8**). The largest of these differences A123^3.27^/I could potentially form steric clashes with the *ortho*-cyclopropylbenzyl group of MIPS3614, consistent with the A123^3.27^I mutant resulting in a modest reduction in EC_50_ shift by MIPS3614 (**Fig. 4e**).

This selectivity pattern is consistent with previous findings for related compounds. The original allosteric modulator BMS-986187 showed 100-fold selectivity for δOR over μOR in a β-arrestin recruitment assay, and an *ortho*-bromobenzyl analogue demonstrated >200-fold selectivity, supporting the importance of the *ortho*-benzyl substitution for receptor discrimination^23^. However, our Nb33 recruitment assay revealed different selectivity profiles. While BMS-986187 showed no allosteric modulation of μOR, MIPS3614 displayed similar activity at both μOR and δOR (**Supplementary Fig. 8c-d**). These results demonstrate that while this allosteric site can confer selectivity between opioid receptor subtypes, the degree of selectivity is inherently limited by the highly conserved nature of the binding site across the opioid receptor family.

### Molecular mechanism of positive allosteric modulation

Along with the A_2A_ adenosine receptor^43^, the δOR and µOR have served as model systems for studying the allosteric effects of sodium ions on class A GPCRs^44,45^. The sodium site is highly conserved across class A GPCRs^34^, including the strongly conserved aspartate at position 2.50 (D^2.50^), and an Na^+^ bound in this pocket stabilizes the inactive receptor conformation. Upon activation, rearrangement of the TM helices leads to a collapse of the sodium pocket and expulsion of the ion, which likely is an essential part of the activation mechanism^34,46^. This process may also be associated with protonation of D^2.50^, disrupting the ionic interaction with the sodium and thereby facilitating receptor activation^46–48^. At δOR, the sodium pocket includes N131^3.35^, at a position that is typically (∼70%) occupied by a hydrophobic residue in GPCRs^33^. Comparison of N131^3.35^ in the inactive (PDB: 4N6H) and active state DADLE-and DADLE-MIPS3614-bound structures reveals a change in the rotamer, resulting in a 5.5 Å movement of the side chain (**Fig. 5a**). In the inactive state, N131^3.35^ points towards the sodium site and interacts directly with the sodium ion, whereas in the active state, the side chain rotates outward from the TM bundle. Notably, the carbonyl oxygen of MIPS3614 forms a hydrogen bond interaction with N131^3.35^ in the cryo-EM structure, which may further stabilize the active state. Attempts to interrogate the contribution of N131^3.35^ by mutagenesis to an alanine or a serine resulted in reduced E_max_ in response to DADLE and surface expression levels (**Fig. 4e-h**), consistent with prior attempts^33^. However, mutation of N131^3.35^ to an aspartic acid did not significantly affect DADLE E_max_ or expression but completely abolished the leftward potentiation of the DADLE response in the presence of MIPS3614 (**Fig. 4i**).

**Fig. 5.**
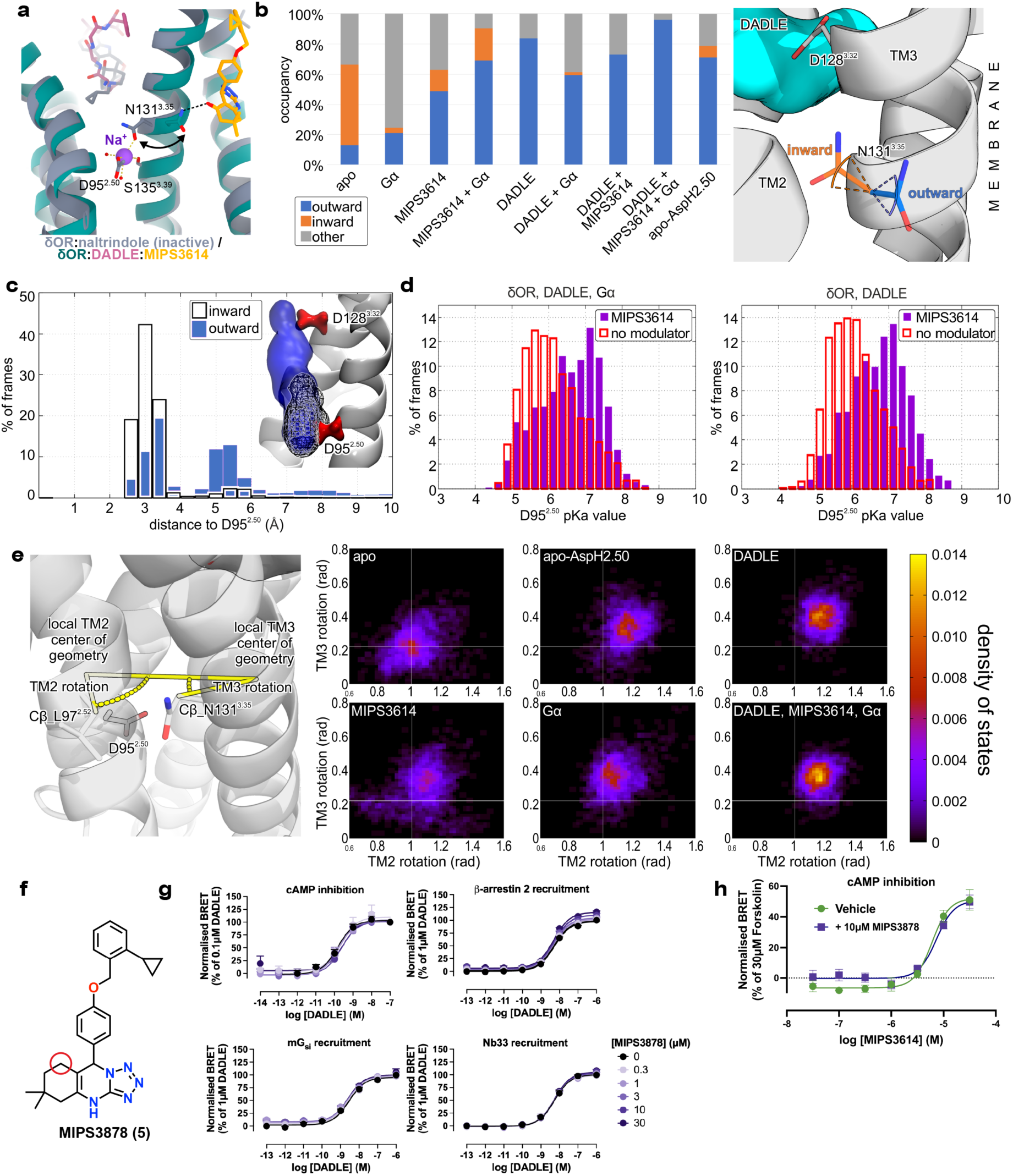
Mechanism of allosteric modulation by disrupting the sodium binding site. **(a)** Structural comparison of the conserved sodium binding site between inactive δOR (δOR:naltrindole, grey; PDB:4N6H) and active MIPS3614-bound δOR (δOR:DADLE:MIPS3614, teal/orange). The overlay shows a 5.5 Å outward conformational change of N131^3.35^ away from the sodium binding site (Na⁺ shown as a purple sphere) in the active state. Key sodium site residues D95^2.50^ and S135^3.39^ are labelled. The carbonyl oxygen from MIPS3614 forms a hydrogen bond with N131^3.35^, stabilizing the outward rotamer conformation associated with receptor activation. (**b**) Occupancy of inward-or outward-pointing rotamers of N131^3.35^ in MD simulations based on the MIPS3614-bound cryo-EM structure. Presence of the PAM, agonist, or Gα protein results in an increase in the occurrence of the outward rotamer compared to the apo state. (**c**) Frequency of D95^2.50^-sodium distances in the simulations of the inactive receptor conformation. The inward conformation of N131^3.35^ stabilizes the sodium ion in the vicinity of D95^2.50^ in nearly all simulation frames (black outlined bars and black mesh). In simulations with N131^3.35^ restrained in the outward conformation, the distribution of distances is altered, with frequent occurrences of sodium near D128^3.32^ in the orthosteric site (blue bars and blue surface). (**d**) Comparison of the p*K*_a_ value of D95^2.50^ calculated for the simulations of the DADLE-bound receptor in the presence (purple bars) or absence (red-outlined bars) of MIPS3614. In the presence of MIPS3614, the p*K*_a_ of D95^2.50^ is shifted towards higher values, favouring protonation of the side chain carboxylate. (**e**) Heatmaps of the distribution of the TM2 and TM3 rotation angle values, measured as depicted on the cartoon receptor representation, from the set of simulations based on the MIPS3614 cryo-EM structure. The presence of PAM, agonist, or Gα protein, as well as D95^2.50^ protonation, induces a concerted rotation of TM2 and TM3. The light-gray lines were added to facilitate comparison between the panels. (**f**) Chemical structure of MIPS3878 (5), a deoxygenated analogue of MIPS3614. The red circle highlights the absence of the carbonyl oxygen in the tricyclic heterocyclic core. (**g**) Pharmacological characterisation of MIPS3878 across multiple signaling pathways showing complete loss of positive allosteric modulation. Data are presented as mean ± SEM with n=4 experiments performed for all assays except cAMP inhibition (n=5). (**h**) A PAM competition assay using MIPS3614 as an allosteric agonist in a cAMP inhibition assay (see Fig. 2c). Addition of 10 µM MIPS3878 did not alter the potency of MIPS3614, suggesting no measurable binding of MIPS3878 at 10 µM in this assay. Data are from n=4 experiments.

To further investigate the influence of the allosteric modulator on the sodium binding site, we carried out molecular dynamics (MD) simulations of the MIPS3614-bound δOR conformation. We first compared the side chain conformation of N131^3.35^ in simulations of the receptor bound to Gα protein, DADLE or MIPS3614 either individually or in combinations, as well as in the apo state (**Fig. 5b**). In agreement with our structural data, the N131^3.35^ side chain almost exclusively remained in an outward conformation in the presence of the agonist, PAM, and G protein. As expected, based on the inactive structure of the δOR, the side chain relaxed to the inward conformation in the simulation of the apo receptor. Notably, DADLE and MIPS3614, either individually or combined, stabilized the outward N131^3.35^ conformation over the inward one. This shows that both agonist and PAM favour an active-like receptor conformation in the sodium pocket region.

To assess how the N131^3.35^ orientation affects sodium binding, we performed MD simulations of the inactive receptor (PDB: 4N6H) in the orientation observed in the X-ray structure and with the asparagine side chain restrained in an outward conformation. The sodium ion present in the inactive structure remains bound at D95^2.50^ in simulations with N131^3.35^ unrestrained. In contrast, the outward rotation of the N131^3.35^ side chain destabilizes the ionic interaction (**Fig. 5c**), and the sodium ion instead moves towards D128^3.32^ in the orthosteric pocket. Furthermore, pK_a_ predictions for D95^2.50^ obtained using PROPKA^49^ on simulation snapshots, show that MIPS3614 binding induces receptor conformations that increase the protonation of the side-chain carboxylate (**Fig. 5d**).

To identify potential large-scale motions in the receptor, principal component analysis (PCA) was applied to the simulations based on the MIPS3614 cryo-EM structure. Subsequently, the motions captured in the first principal components were analysed in the MD trajectories. This revealed that MIPS3614, DADLE, or Gα protein binding, as well as D95^2.50^ protonation, each induce bending of TM3 that is consistent with available active and inactive δOR structures (**Supplementary Fig. 9**). Moreover, this movement is associated with a concerted rotation of TM2 and TM3 compared to the apo state. In the simulation with the receptor bound to Gα protein, MIPS3614 and DADLE, the mangle-like TM2/TM3 rotation leads to high occupancy of the outward conformation of N131^3.35^ that enables hydrogen bonding to MIPS3614. The receptor conformations stabilized by the MIPS3614 hence contribute to displacing the sodium ion and favour protonation of D95^2.50^, facilitating receptor activation.

To further validate the mechanism, we synthesised an analogue lacking the carbonyl oxygen on the tetrazoloquinazolinone core that forms the critical hydrogen bond to N131^3.35^ (**Fig. 5f**). This deoxygenated analogue, MIPS3878 (**5**), was designed to test the functional importance of this specific interaction. MIPS3878 showed a complete loss of modulatory activity across multiple functional assays (**Fig. 5g**). In the Nb33 recruitment assay, MIPS3878 produced no shift in the EC_50_ of DADLE, and similar results were observed in cAMP inhibition, β-arrestin2 recruitment, and mG_si_ coupling. To determine whether MIPS3878 was acting as a neutral allosteric ligand (NAL), binding to the allosteric site without modulating it, or had completely lost binding affinity, we performed competition studies. We assessed increasing concentrations of MIPS3614 in cAMP inhibition assays conducted in agonist mode (absence of orthosteric ligand), both with and without 10 μM MIPS3878 (**Fig. 5h**). In this experimental design, a neutral allosteric ligand (NAL) would function as an allosteric antagonist, competing with the response elicited by MIPS3614. However, MIPS3878 did not affect the response to MIPS3614 in cAMP assays at concentrations up to 10 µM, suggesting removal of the carbonyl oxygen reduces binding to the allosteric site.

Intriguingly, we observed a slight, but statistically significant concentration-dependent increase in the E_max_ of DADLE with increasing concentrations of MIPS3878 in the β-arrestin 2 assay (**Fig. 5g**). Prior mutation of residue N131^3.35^ to alanine at δOR resulted in constitutive β-arrestin activation^33^. We speculate that the loss of the hydrogen bond with N131^3.35^ may confer biased allosteric properties to MIPS3878, potentially acting as a pathway-selective positive allosteric modulator favouring β-arrestin 2 activation over G protein coupling. This contrasts with BMS-986187, which is biased towards G protein signaling relative to β-arrestin 2 activation^50^, and highlights the importance of Na^+^ binding site in controlling the active conformation of class A GPCRs^51^.

### Structure-guided optimisation of 8OR allosteric modulators

The initial structure-activity relationship (SAR) studies for the BMS-based scaffold focused on modifications to the substituent in the 9-position^52^, while a subsequent investigation examined modifications to the tricyclic heterocyclic core^32^. Based on the binding pose of MIPS3614 in our δOR structure, we hypothesized that extending substituents from the 4-position of the heterocyclic core would enhance interactions with residues within the allosteric pocket, particularly S177^4.54^.

A new series of compounds was synthesized (**Fig. 6a, 6a-i**) and evaluated for their ability to potentiate DADLE response in the Nb33 recruitment assay (**Fig. 6b-c**). The new analogues generally demonstrated 10-fold improved binding affinity (p*K*_B_) compared to MIPS3614 (**Fig. 6b**). However, compounds containing polar substituents, specifically the ethanol and acetic acid groups in **6c** and **6e**, showed significantly reduced functional cooperativity and binding affinity. This loss of activity was attributed to unfavourable polar interactions within the predominantly hydrophobic allosteric pocket. Based on these results, we focused subsequent studies on MIPS3983 (**6f**), which contains a methyl acetate substituent and demonstrated improved binding affinity (p*K*_B_ = 5.4 compared to 4.6 for MIPS3614) while retaining a substantial cooperativity factor of 12. The methyl acetate group appeared to provide an optimal balance between enhanced binding through additional interactions and maintenance of appropriate physicochemical properties for the binding site. We choose MIPS3983 for structural studies because of the larger size of the methyl acetate group compared to appendages of compounds **6a**, **6b**, **6d**, and **6g**, which had comparable p*K*_B_ and *αβ* values to MIPS3983.

**Fig. 6.**
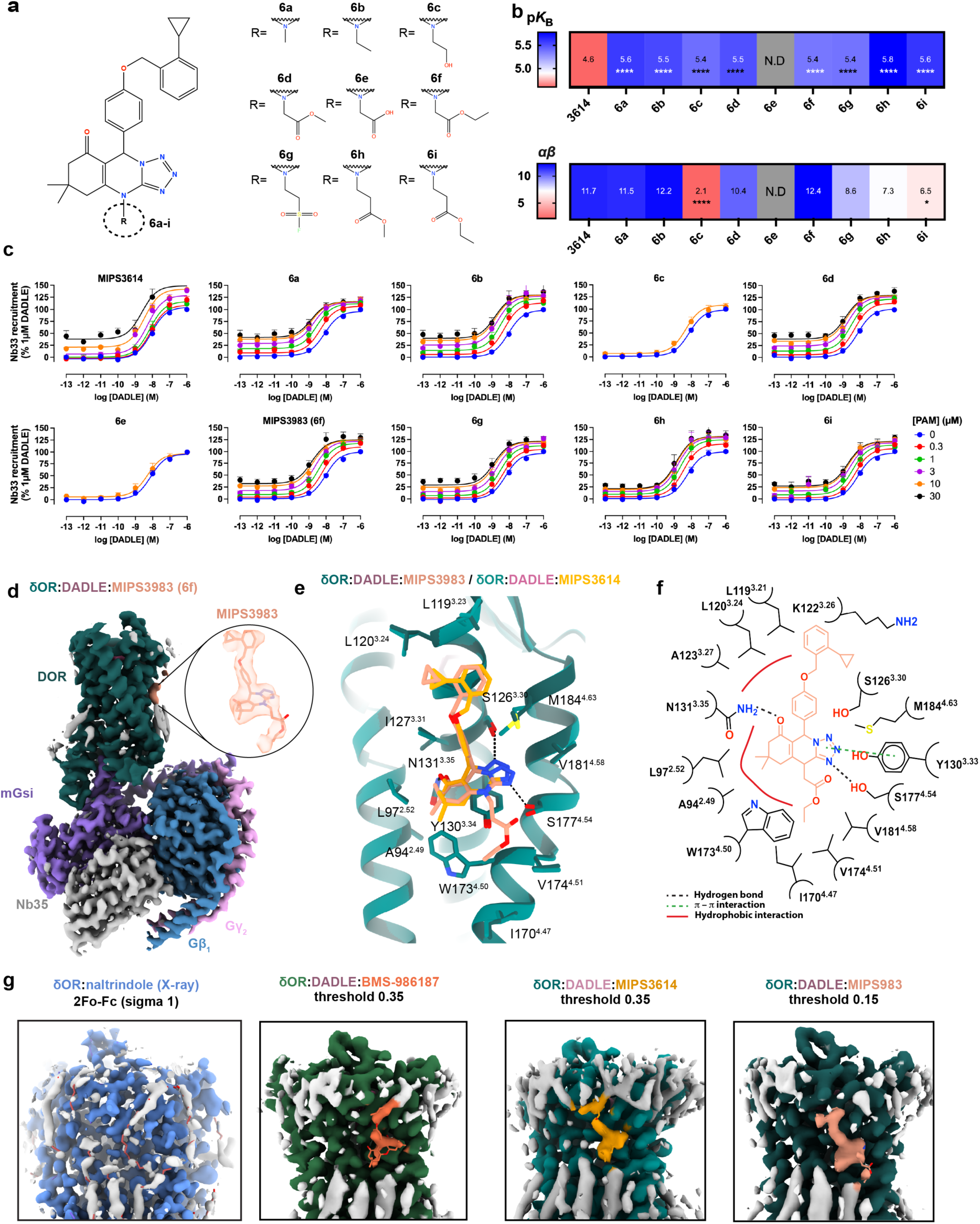
Structure-guided optimization of δOR positive allosteric modulators. **(a)** Chemical scaffold showing the tricyclic heterocyclic core structure with variable R-group substituents at the secondary amine position (dashed circle). Nine analogues were synthesized (6a-i) to explore structure-activity relationships and enhance interactions with the allosteric binding pocket, particularly targeting residue S177^4.54^. (**b**) Heat maps summarising the pharmacological properties of the new analogues derived from concentration response curves in the Nb33 recruitment assay (shown in **c**). Top panel shows binding affinity (p*K*_B_) values, and the bottom panel shows functional cooperativity (*αβ*) values with mean values overlaid on the map. N.D. = not determined. Statistical significance was determined by one-way ANOVA with a Dunnett’s post-test using MIPS3614 for comparison (*P=0.0304; ****p < 0.0001). Data are from n=7 experiments for MIPS3614, n=6 for **6a**, n=4 for **6b**, n=7 for **6c**, n=5 for **6d**, n=5 for **6e**, n=5 for **6f,** n=5 for **6g**, n=4 for **6h,** n=6 for **6i**. (**d**) Cryo-EM structure of the δOR-DADLE-MIPS3983 complex showing improved cryo-EM density in the allosteric binding site. The inset displays cryo-EM density from the receptor-focused map at a contour of 0.15. (**e**) Detailed view of MIPS3983 binding interactions in the allosteric pocket. Compared to MIPS3614, the tricyclic heterocyclic core of MIPS3983 appears to rotate in a manner that allows hydrogen bonding to both S126^3.30^ and S177^4.54^. (**f**) Molecular interaction diagram of MIPS3983 with δOR residues. Hydrogen bonds are denoted by black dashed lines, π-stacking interactions with green dashed lines, and hydrophobic interactions with solid red lines. (**g**) Comparison of density maps across different δOR structures highlighting potential lipid/cholesterol binding sites near the MIPS3614 allosteric site.

Using the same methodology employed for MIPS3614, we determined a cryo-EM structure of the human δOR in complex with DADLE and MIPS3983 (**Fig. 6d, Supplementary Fig. 10 and Table 1**). MIPS3983 was bound to the same allosteric site as MIPS3614, with notably improved interpretability of the cryo-EM density around the tricyclic heterocyclic core and the *ortho*-cyclopropylbenzyl group. This enhanced density definition likely reflects the improved binding affinity of MIPS3983. The additional methyl acetate substituent showed more ambiguous density, suggesting either conformational flexibility or potential steric interactions with surrounding detergent or cholesterol molecules observed near the binding site. Nevertheless, the methyl acetate group could be modelled into the cryo-EM density extending below the primary allosteric binding pocket. The tricyclic heterocyclic core and 9-(4-((*ortho*-cyclopropylbenzyl)oxy)phenyl) groups of MIPS3983 formed the same key interactions observed with MIPS3614. Additionally, MIPS3983 established a new hydrogen bond between S177^4.54^ and the nitrogen atom of the heterocyclic core, confirming our design hypothesis (**Fig. 6d**). The methyl acetate substituent extended downward from the heterocyclic core, forming additional hydrophobic interactions with W173^4.50^ and V174^4.51^.

These structural and functional studies demonstrate that the allosteric pocket can accommodate additional substituents that enhance binding affinity through new productive interactions. The success of MIPS3983 validates the structure-based design approach and provides a foundation for further optimization of this allosteric modulator scaffold. The improved affinity and retained cooperativity of MIPS3983 make it a valuable tool compound for studying δOR allosteric modulation.

## Discussion

Multiple approaches have been explored in the search for safer opioid analgesics, including partial agonists that provide submaximal receptor activation^53^, biased agonists that preferentially activate G protein over β-arrestin pathways^54^, and allosteric modulators that enhance endogenous ligand activity^30^. Our structural and functional characterisation of δOR PAMs provides the first molecular framework for understanding how allosteric modulation can be achieved at δOR, opening new avenues for therapeutic development.

Although we could not confidently model BMS-986187 binding due to insufficient cryo-EM density, we successfully determined the allosteric binding site using MIPS3614, a more soluble analogue with similar pharmacological properties. The shared loss of allosteric activity at the M184^4.61^F mutant confirms that both compounds bind the same site. Our structural determination corrects previous computational predictions of the BMS-986187 binding site, which had relied on molecular docking and limited mutagenesis data^55^. The experimentally determined binding site reveals interactions and binding mechanisms that were not anticipated from earlier modelling efforts, highlighting the critical value of high-resolution structural data for understanding allosteric modulation.

The δOR allosteric binding site was located in a lipid-facing pocket formed by residues from transmembrane helices 2, 3, and 4. Lipid-facing allosteric sites are emerging as a common feature of GPCR-positive allosteric modulators, though with distinct binding mechanisms^56^. The CB₁ receptor PAM ZCZ011 binds at a similar transmembrane location^57^ but utilises purely hydrophobic interactions without engaging the sodium binding site (**Supplementary Fig. 11a-d**). In contrast, the μOR-selective PAM BMS-986122 occupies a different lipid-facing pocket 15 Å away^58^, formed by TMs 3, 4, and 5 (**Supplementary Fig. 11e-h**). The selectivity of BMS-986122 for μOR over δOR likely stems from a steric clash with the larger methionine residue at position 3.45 in δOR versus threonine in μOR.

Our identification of the δOR PAM binding site reveals a novel mechanism whereby allosteric modulation is achieved through direct stabilisation of the sodium binding site in its collapsed, active-state conformation^34^. Prior studies have suggested that the mechanism of action for opioid receptor PAMs BMS-986122 and BMS-986187 involves destabilising the inactive Na^+^-bound conformation of the receptors, with PAM binding demonstrating a rightward shift in NaCl concentration response curve^59,60^. These pharmacological findings, along with allosteric competition assays, provided evidence of a potential conserved OR allosteric site. However, in light of our δOR-MIPS3614 structure and the µOR-BMS-986122 structure, it is clear the allosteric binding sites are distinct, yet conformationally linked, explaining the observed competition between BMS-986122 and BMS-986187^59^. This effect of PAM binding on NaCl concentration response curves is likely common to class A PAMs that stabilize the active conformation of GPCRs.

Our MD simulations show that the interaction between MIPS3614 and N131^3.35^ contributes to the displacement of the sodium ion and favours the protonation of D95^2.50^, thereby facilitating receptor activation. The simulations reveal concerted motions of TM2 and TM3 that impact the sodium pocket architecture. These conformational rearrangements alter the local environment surrounding the conserved D95^2.50^, shifting its p*K*_a_ toward higher values and thus favouring protonation. This protonation event further destabilizes sodium binding and promotes ion expulsion, creating a positive feedback mechanism that stabilizes the receptor’s active conformation. The coupling between MIPS3614 binding, N131^3.35^ reorientation, and sodium site disruption represents a key mechanistic link between allosteric modulation and receptor activation.

The high resolution of our cryo-EM maps (**Fig. 6g**) enabled identification of lipid binding sites both above and below the δOR allosteric pocket, highlighting the role of membrane lipids in defining the architecture of allosteric binding sites^56,61^. These lipid interactions help explain structure-activity relationships observed in medicinal chemistry campaigns. For example, SAR studies of the BMS-986187 tolyl group showed that polar heterocycles and larger aromatic rings reduced PAM activity^52^, suggesting size constraints and the importance of hydrophobic interactions with nearby lipids. Similarly, MIPS3614 analogues with bulky modifications to the tetrazoloquinazolinone secondary amine (**6c**/**6e**) showed reduced activity, likely due to steric clashes with lipids positioned below the binding site. These findings demonstrate that membrane lipids are integral components of the allosteric binding site and must be considered in structure-based drug design efforts^7^.

Our findings also present new opportunities for designing novel classes of allosteric ligands. The identification of lipid-binding sites adjacent to the allosteric pocket suggests that tethering allosteric modulators to membrane-targeting moieties could enhance binding affinity and selectivity. This approach has proven successful for other GPCRs, such as neurokinin 1 receptor antagonists linked to lipid groups that show enhanced potency, affinity, and duration of action^62^. Additionally, given recent findings that δOR activation from endosomes provides sustained pain inhibition^7^, lipid-conjugated PAMs might facilitate subcellular targeting to specific membrane compartments, potentially extending therapeutic effects.

From a therapeutic perspective, δOR PAMs offer several potential advantages over direct agonists, including preservation of endogenous signaling patterns, which could reduce the propensity for tolerance development^27^. Although this study further validates the potential of utilising δOR PAMs to treat GI disorders^29^, future studies will be necessary to validate the potential of δOR PAMs for treating pain. Our detailed structural and mechanistic insights establish a robust platform for structure-based drug design of improved δOR PAMs. Medicinal chemistry efforts can now target the subtle sequence differences identified between opioid receptor subtypes or exploit the lipid-binding regions to develop next-generation compounds with enhanced selectivity and therapeutic efficacy.

## Acknowledgements

This work was funded by a National Health and Medical Research Council (NHMRC) Investigator Grant 1196951 (D.M.T.) and 2026533 (A.B.G), an NHMRC Ideas Grant 2021675 (D.P.P., S.E.C., C.V., J.I.M.), an NHMRC Program Grant 1150083 (A.C.), an Australian Research Council (ARC) DECRA DE240100931 (A.B.G), a Sven and Lilly Lawski Foundation, stipend no. N2024-0043 (D.B.), and a Swedish Research Council Grant 2021-4186 (J.C.) The National Academic Infrastructure for Supercomputing in Sweden (NAISS), partially funded by the Swedish Research Council through grant agreement no. 2022-06725. The work was supported by the Monash University Ramaciotti Centre for Cryo-Electron Microscopy, and the Monash eResearch capabilities, including M3 High-performance computing. Figures were created with UCSF Chimera and ChimeraX, developed by the Resource for Biocomputing, Visualization, and Informatics at the University of California, San Francisco, with support from National Institutes of Health R01-GM129325 and the Office of Cyber Infrastructure and Computational Biology, National Institute of Allergy and Infectious Diseases, as well as with VMD and PyMOL.

## Author contributions

A.B.G, P.J.S., C.V., D.M.T. designed the overall project. J.I.M, O.D. designed, expressed, and purified protein samples. J.I.M., H.V. performed sample vitrification and cryo-EM imaging. J.I.M. and D.M.T. processed the EM data and generated and analysed atomic models. M.N., V.P., N.B., A.B.G. performed pharmacology experiments in recombinant systems. S.A. performed pharmacology experiments in native tissues and whole organ experiments. O.D., P.J.S, B.C., M.J. designed compounds. O.D. synthesised and characterised compounds. D.B. performed and analysed molecular dynamics simulations. M.N., V.P., N.B., A.B.G., S.A., D.P.P, S.E.C., C.V., D.M.T analysed pharmacology data. D.P.P., S.E.C., A.C., B.C., M.J., J.C., A.B.G, P.J.S., C.V., D.M.T. provided project supervision. J.I.M, M.N, O.D., A.B.G, D.M.T. wrote the manuscript with contributions from all authors.

## Methods

### Structure

#### Expression and purification of Nb35

Nb35 was expressed using an autoinduction method in BL21 (DE3) Rosetta *Escherichia coli*^63,64^. Briefly, cells were transformed and grown at 37 °C under antibiotic selection with 100 μg/mL carbenicillin and 35 μg/mL chloramphenicol in autoinduction media (50 mM Phosphate buffer, pH 7.2, 2% tryptone, 0.5% yeast extract, 0.5% NaCl, 0.6% glycerol, 0.05% glucose and 0.2% lactose). At an OD_600_ of 0.7, the cells were transferred to 20 °C for approximately 16 h before being harvested by centrifugation. Cell pellets were stored at -80 °C until purification. Nb35 was purified using Ni-affinity chromatography as previously described^65^.

#### Expression and purification of δOR-mG_si_-DADLE complex

The full-length human delta opioid receptor gene was modified to include an N-terminal hemagglutinin (HA) signal sequence and FLAG-tag, and a C-terminal 8xHis tag, GGGS linker, 3C protease cleavage site, and mini-G_si_ (mG_si_)^66,67^. The resulting FLAG-δOR-mG_si_ construct was cloned into the pFastBac vector (Invitrogen) for insect cell expression.

FLAG-δOR-mG_si_ and Gβ_1_γ_2_ were co-expressed in Tni insect cells (Expression Systems), using the Bac-to-bac baculovirus expression system (Invitrogen). The Tni cells were grown to a density of 4×10^6^ cells per ml in ESF 291 serum-free media (Expression Systems) and infected with a 4:1 ratio of FLAG-δOR-mG_si_:Gβ_1_γ_2_ baculoviruses. Cells were incubated for 48-60 h at 27 °C, before harvesting by centrifugation (6,000 g, 20 min, 4 °C). and cell pellets stored at - 80 °C.

For the purification of δOR-mG_si_-DADLE complex, the cell pellet was thawed and resuspended in lysis buffer (20 mM HEPES, pH 7.4, 5 mM MgCl_2_) with protease inhibitors (0.2 mM PMSF, 5 µg/mL leupeptin, 5 µg/mL soybean trypsin inhibitor and 1 mM benzamidine hydrochloride), 500 U benzonase and 1 μM DADLE peptide. The resuspension was stirred at room temperature for 15 min to allow for lysis before centrifugation at 30,000 g for 20 min at 4 °C. The pellet was resuspended in solubilization buffer (20 mM HEPES, pH 7.4, 100 mM NaCl, 5 mM MgCl_2_, 5 mM CaCl_2_, 0.5% LMNG, 0.03% CHS) with protease inhibitors (0.2 mM PMSF, 5 µg/mL leupeptin, 5 µg/mL soybean trypsin inhibitor and 1 mM benzamidine hydrochloride), 500 U benzonase. The resuspended pellet was homogenised with a Dounce and complex formation initiated with the addition of 1 μM DADLE peptide, 1 mg Nb35 and 25 mU/mL apyrase. The complex was incubated with stirring for 2 h at 4 °C before centrifugation at 30,000 g for 30 min at 4 °C to remove insoluble material. The complex was then bound to M1 anti-FLAG-affinity resin by gravity flow before washing with wash buffer (20 mM HEPES, pH 7.4, 100 mM NaCl, 5 mM MgCl_2_, 5 mM CaCl_2_, 0.01% LMNG, 0.0006% CHS and 1 μM DADLE). The complex was eluted with elution buffer (20 mM HEPES, pH 7.4, 100 mM NaCl, 5 mM MgCl_2_, 0.01% LMNG, 0.0006% CHS, 1 μM DADLE, 10 mM EGTA and 0.1 mg/ml FLAG peptide) before concentration in an Amicon Ultra-15 100 kDa molecular mass cut-off centrifugal filter unit (Millipore). The complex was further purified by size exclusion chromatography on a Superdex 200 Increase 10/300 GL (Cytiva) in 20 mM HEPES, pH 7.4, 100 mM NaCl, 5 mM MgCl_2_, 0.01% LMNG, 0.0006% CHS and 1 μM DADLE. Final fractions were assessed by SDS-PAGE, pooled, concentrated to 10 mg/mL, and flash-frozen in liquid nitrogen before storage at -80 °C

#### Grid preparation and imaging

For the allosteric complexes, the sample was thawed and the allosteric ligand was added (100 μM for BMS-986187, 200 μM for MIPS3614, 100 μM for MIPS3983) and incubated overnight on ice at 4 °C. No allosteric ligand was added for the 8OR-mG_si_-DADLE complex structure. 3 µL of sample was applied to glow-discharged (15 mA, 180 s) UltrAufoil R1.2/1.3 300 mesh holey grid (Quantifoil) and was frozen in liquid ethane using a Vitrobot mark IV (Thermo Fisher Scientific) at 100% humidity and 4°C with a blot time of 2 s and blot force of 10 – 14. Data were collected using a G1 Titan Krios microscope (Thermo Fisher Scientific) equipped with S-FEG, a BioQuantum energy filter and K3 detector (Gatan). The Krios was operated at an accelerating voltage of 300 kV with a 50 μm C2 aperture, 100 μm objective aperture inserted, and zero-loss filtering (10 eV slit width), at 105 kX magnification in nanoprobe EFTEM mode. Data were collected using aberration-free image shift (AFIS) with Thermo Fisher EPU software.

#### Cryo-EM data processing

All datasets were collected at 0.82 Å/pix and processed using a standardised workflow. Movies were motion corrected using UCSF MotionCor2^68^ and CTF parameters estimated using GCTF^69^. Movies for δOR-mG_si_-DADLE-MIPS3983 were motion corrected and output in float16 using Relion v5.0^70,71^ and CTF parameters were estimated using CTFFIND 4.1^72^. Particles were picked using the general model of crYOLO^73^ and extracted with RELION v3.1^70,71^ (v5.0 for δOR-mG_si_-DADLE-MIPS3983) at 60 pix, 4.92 Å/pix. Following import to cryoSPARC v4.1.2^74^ for 2D classification and Ab-Initio model generation, heterogeneous refinement was performed with 2 initial models and a junk trap (1 initial model for δOR-mG_si_-DADLE-MIPS3983). Full-sized particles were reextracted at 360 pix, 0.82 Å/pix and subjected to subsequent rounds of 2D classification and heterogeneous refinement in cryoSPARC v4.1.2. Selected particles underwent Bayesian Polishing in RELION and final non-uniform refinement with CTF refinement. For δOR-mG_si_-DADLE-MIPS3983, an additional 3D classification step without alignment was performed (6 classes, T=10) before final refinement. Dataset-specific details can be found in Supplementary Fig. 2, 5, 6, and 10.

#### Model building and refinement

An initial δOR template model was generated from the cryo-EM structure of the µ opioid receptor (PDB: 6DDE)^75^. An initial model for the miniG_si_ was generated from the structure of CCK1:mG_sqi_ complex (PDB: 7MBY) and the Gβ_1_Gγ_2_, and Nb35 were taken from the CCK1:DNG_s_ complex (PDB: 7MBX)^40^. Models were placed into the cryo-EM maps using UCSF ChimeraX^76^ and rigid-body-fit using PHENIX^77^. Models were refined with iterative rounds of manual model building in Coot^78^ and ISOLDE^79^, and real-space refinement in PHENIX. Restraints for all ligands were generated using the GRADE web server (https://grade.globalphasing.org). Model validation was performed with MolProbity^80^ and the wwPDB validation server^81^. Figures were generated using UCSF ChimeraX^76^ and PyMOL (Schrödinger).

#### Computational chemistry

MD simulations were performed using Gromacs 2024.2^82^ patched with Plumed 2.9.2. Amber03, GAFF2 and Slipids force fields were used for protein, small molecule ligands and membrane, respectively^83–86^. The simulation box with the receptor and an asymmetric membrane was built using CHARMM-GUI^87^. The cryo-EM structure reported in this manuscript was used for the active-state receptor simulations, and the 4N6H PDB structure was used for the inactive-state calculations. Receptor termini were capped with *N*-methyl and acetate caps. The membrane was composed of nine lipid types and included cholesterol, sphingomyelin, phospholipid headgroups: phosphatidylcholine, phosphatidylethanolamine, phosphatidylserine, phosphatidylglycerol, and palmitoyl, oleyl and linoleoyl lipid tails. The simulation box was filled with TIP3P water and 0.15 M NaCl. Ligand partial charges were calculated with PyRED^88^, and ligand topologies were generated using ACPYPE^89^. The target volume was achieved with 5 ns NPT simulation with the Berendsen barostat, followed by 300 ns box pre-equilibration in the NVT ensemble with position restraints of 10,000 kJ/mol nm^2^ imposed on the protein. After the membrane pre-equilibration, the simulation boxes with specified ligand sets were prepared and equilibrated. Insertion of MIPS3614 required removal of the overlapping lipids and was followed by an additional 250 ns NVT equilibration with maintained restraints on protein atoms. Energy minimization of each system was then performed until the maximal force was less than 100.0 kJ/mol/nm. This was followed by 500 ps of equilibration run in the NVT ensemble and 1 ns in NPT ensemble with Parrinello-Rahman barostat and position restraints of 1000 kJ/mol nm^2^ applied to the protein main chain and Cα atoms, respectively. The dimensions of the simulation boxes were approximately 83 Å x 83 Å x 141 Å with Gα protein and 83 Å x 83 Å x 107 Å without Gα protein. The Nose-Hoover thermostat was used for temperature coupling at 309 K, and the Parrinello-Rahman barostat was used for semiisotropic pressure coupling at 1 bar for the production runs. Long-range electrostatics were treated with the particle mesh Ewald method^90^. LINCS algorithm^91^ was used to constrain the hydrogen-heavy atom bonds, and a time step 2 fs was used. All-atom unbiased MD production simulations were run for 1 μs in six replicas per complex, and trajectory frames were saved every 400 ps. To perform PCA of all complexes in a common subspace, trajectories were truncated to the receptor heavy atoms using gmx trjconv, concatenated and fitted using the Cα atoms. PCA of the resultant trajectory was then performed with the Gromacs tools gmx covar and gmx anaeig. Trajectory analyses were performed with Gromacs tools, the open-source, community-developed PLUMED library (version 2.9.2)^83^, and VolMap tool. PROPKA was used to calculate p*K*_a_ values^49^. Plots and heat maps were generated using Gnuplot^49^. Molecular visualizations were created with PyMOL^92^ and VMD^93^.

#### Enzyme-linked immunosorbent assay (ELISA)

Flp-In-HEK293 cells were seeded at 50,000 cells per well into poly-D-lysine–coated 48-well plates, and 4-6 h later transfected with 100 ng/well of FLAG-hδOR WT or mutants using a 1:6 ratio of DNA:PEI in 150 mM NaCl. Cells were fixed with 3.7% (v/v) paraformaldehyde in tris-buffered saline (TBS) for 30 min. For receptor expression, cells were permeabilized by 30-min incubation with 0.5% (v/v) NP-40 in TBS. Cells were then incubated in blocking buffer [1% (w/v) skim milk powder in 0.1 M NaHCO_3_ for 4 h at room temperature and incubated with mouse M2 anti-FLAG antibody (Sigma, #F1804, 1:2000, 2 hours at 4°C). After washing three times with TBS, cells were incubated with anti-mouse IgG, HRP-linked antibody (Cell Signaling Technology, Cat#7076, 1:2000) for 1 h at room temperature. Cells were washed and stained using TMB substrate (Sigma-Aldrich). Absorbance at 450 nm was measured using an EnVision Multilabel Reader (PerkinElmer).

#### Functional assays

For cAMP inhibition, FlpIn HEK293 cells stably expressing human 8OR were seeded in 10 cm dishes and transfected with 5 μg of cAMP sensor using YFP-Epac-RLuc (CAMYEL) BRET sensor. For Nb33, mG_si_, or β-arrestin 2 recruitment assays, FlpIn HEK293 cells were seeded in 10 cm dishes and transiently transfected with 1 μg of donor (human δOR-RLuc8 WT or mutant) and 4 μg of acceptor (nanobody33-Venus, miniG_si_-Venus or β-arrestin2-YFP) using linear polyethylenimine (Polysciences, Warrington, PA, USA) at a DNA:PEI ratio of 1:6 in 150 mM NaCl. Twenty-four h post-transfection, 30,000 cells/well were seeded in poly-D-lysine-coated 96-well white CulturPlates (PerkinElmer, Glen Waverly, VIC, Australia) and incubated for twenty-four h at 37 °C and 5% CO_2_. Forty-eight hours post-transfection, the cells were washed and then incubated with Hanks’ balanced salt buffer containing 10 mM HEPES, pH 7.4, for 30 min at 37 °C. Increasing concentrations of DADLE in the absence or presence of increasing concentrations of each allosteric modulator were added and incubated for 5 min. Cells were then incubated for 7 min with a final concentration of 5 μM coelenterazine H (Nanolight Technology, AZ) and for cAMP inhibition also 30 μM forskolin (Sigma-Aldrich, Castle Hill, NSW, Australia). BRET was measured on a LUMIstar Omega instrument (BMG LabTech, Mornington, VIC, Australia) at 12 min post agonist addition using 475 ± 30 nm/535 ± 30 nm filters. To determine BRET signal, the ratio of light emitted at 535 ± 30 nm by YFP over the light emitted at 475 ± 30 nm by Renilla Luciferase was quantified.

#### Animals

C57Bl/6J mice (6-8 weeks, male) were obtained from the Monash Animal Research Platform. Mice were housed under a 12 h light/dark cycle, with controlled temperature (24 °C) and humidity, and free access to food and water. All procedures involving animals were approved by the Monash Institute of Pharmaceutical Sciences Animal Ethics Committee (approval 40039) and adhered to the ARRIVE guidelines. Mice were humanely killed by cervical dislocation, and the distal colon was removed and placed in modified Krebs buffer (in mM: 118 NaCl, 4.7 KCl, 1.1 MgSO_4_•7H_2_O, 1.18 KH_2_PO_4_, 25 NaHCO_3_, 11.6 glucose, and 2.5 CaCl_2_).

#### Tissue contraction assay

Segments of distal colon were placed under a resting tension of 0.5g and equilibrated for 30 min before use. Neurogenic contractions of the circular muscle were evoked by transmural electrical field stimulation (EFS; 0.5 ms duration, 3 pulses/s, 60V). Once baseline responses to EFS were established, increasing concentrations of Leu-Enk (final concentration 1 pM-10 µM) were added to the organ baths, along with MIPS3614 (1 or 10 µM), BMS-986187 (10 µM) or vehicle (DMSO). EFS was repeated 3 times per test concentration. The peak amplitude of each EFS-evoked contraction was normalised to the baseline response in the absence of test compounds. To confirm tissue viability, carbamoylcholine (carbachol, 10 μM) was applied after each experiment^94^.

#### Spatiotemporal mapping

Propagating contractions were assessed in the intact colon by video mapping, as described in detail^15^. After an equilibration period, motor patterns were recorded with elevated intraluminal pressure for 20 min. The effects of MIPS3614 on the number, velocity, and interval between pressure-evoked colonic motor complexes (CMCs) were assessed relative to vehicle control.

#### Data analysis

GraphPad Prism software (v. 9.0) was used to analyse all functional data and for statistical analysis. Data points are presented as mean ± standard error of the mean (s.e.m.) based on at least three biologically independent experiments in duplicate. Concentration-response curves were fitted using the three-parameter log(agonist) vs. response equation. p*EC*_50_ values were extracted from the curve fit of each individual experiment. *E*_max_ was calculated by subtracting the Bottom from the Top parameter from the curve fit of each individual experiment. Statistical analysis of Δp*EC_50_* and *E*_max_ values was performed using one-way analysis of variance (ANOVA) with Dunnett’s multiple comparison–corrected post hoc test against 8OR WT, with a p-value of < 0.05 considered significant.

When possible, the data were also fit to the operational model of allosterism^42^ to determine equilibrium dissociation constants of agonist and allosteric modulator are represented by *K*_A_ and *K*_B,_ respectively. The efficacy of orthosteric and allosteric ligands was respectively denoted by *τ*_A_ and *τ*_B_. *αβ* represents the cooperativity factor with *α* denoting the cooperativity between agonist and allosteric modulator at the level of binding (binding cooperativity), and *β* representing the cooperativity between agonist and allosteric modulator at the level of efficacy (efficacy cooperativity; or scaling factor). p*K*_B_ and log *αβ* values were extracted from the global fit of the grouped data of at least three biologically independent experiments. Statistical analysis of p*K*_B_ and log *αβ* values was performed using one-way analysis of variance (ANOVA) with Dunnett’s multiple comparison–corrected post hoc test against MIPS3614, with a p-value of < 0.05 considered significant.

#### Chemistry

The stepwise pathway to attain the desired tetrazoloquinazolinone 4 began with the *O*-alkylation of commercially available 4-hydroxybenzaldehyde 1 with 2-bromobenzyl bromide, forming the (bromobenzyloxy)benzaldehyde precursor 2. The aryl bromide 2 subsequently underwent a Suzuki coupling with cyclopropylboronic acid utilizing palladium(II) acetate in the presence of tricyclohexylphosphine to yield the (cyclopropylbenzyloxy)benzaldehyde intermediate 3. Formation of the tetrazoloquinazolinone 4 was achieved through a one-pot intramolecular cyclocondensation of benzaldehyde 3 with an equivalent of each 5,5-dimethylcyclohexane-1,3-dione and 5-aminotetrazole in MeOH with catalytic quantities of conc. H_2_SO_4_ to aid the intramolecular cyclocondensation. The racemic mixture of 4 was further separated into its corresponding enantiomers (*R*)-4a and (*S*)-4b through preparative chiral HPLC of 4 utilizing cellulose-2 column packing under normal-phase conditions (10% PET/EtOH). LiAlH_4_ was effective in reducing the tetrazoloquinazolinone carbonyl of 4 to yield the tetrazoloquinazoline 5. The complete reduction of the carbonyl moiety (as opposed to partial reduction to the corresponding alcohol) appeared to be a feature of the carbonyl species being in conjugation with the secondary amine, with similar outcomes being cited in literature^95^. With the tetrazoloquinazolinone 4 synthesized, we were able to further derivatize from the secondary amine of the core to form the *N*-substituted analogs 6a-i. The *N*-substituted analogs 6a-d,f were formed through simple nucleophilic substitution reactions with the appropriate alkyl halide, whereas the extended chain analogs 6g-i were generated *via* Michael addition reactions to incorporate the additional methylene. A caveat to forming the *N*-substituted analogs was the long reaction times and poor conversion to product, which was a result of the nature of the secondary amine being in conjugation to both the adjacent tetrazole and *α*,*β*-unsaturated ketone, minimizing the nucleophilicity of the amine. Base-catalyzed ester hydrolysis of the methyl ester 6d using LiOH.H_2_O in a H_2_O/THF system furnished the corresponding acetic acid analog 6e in quantitative yield.

**Scheme 1.**
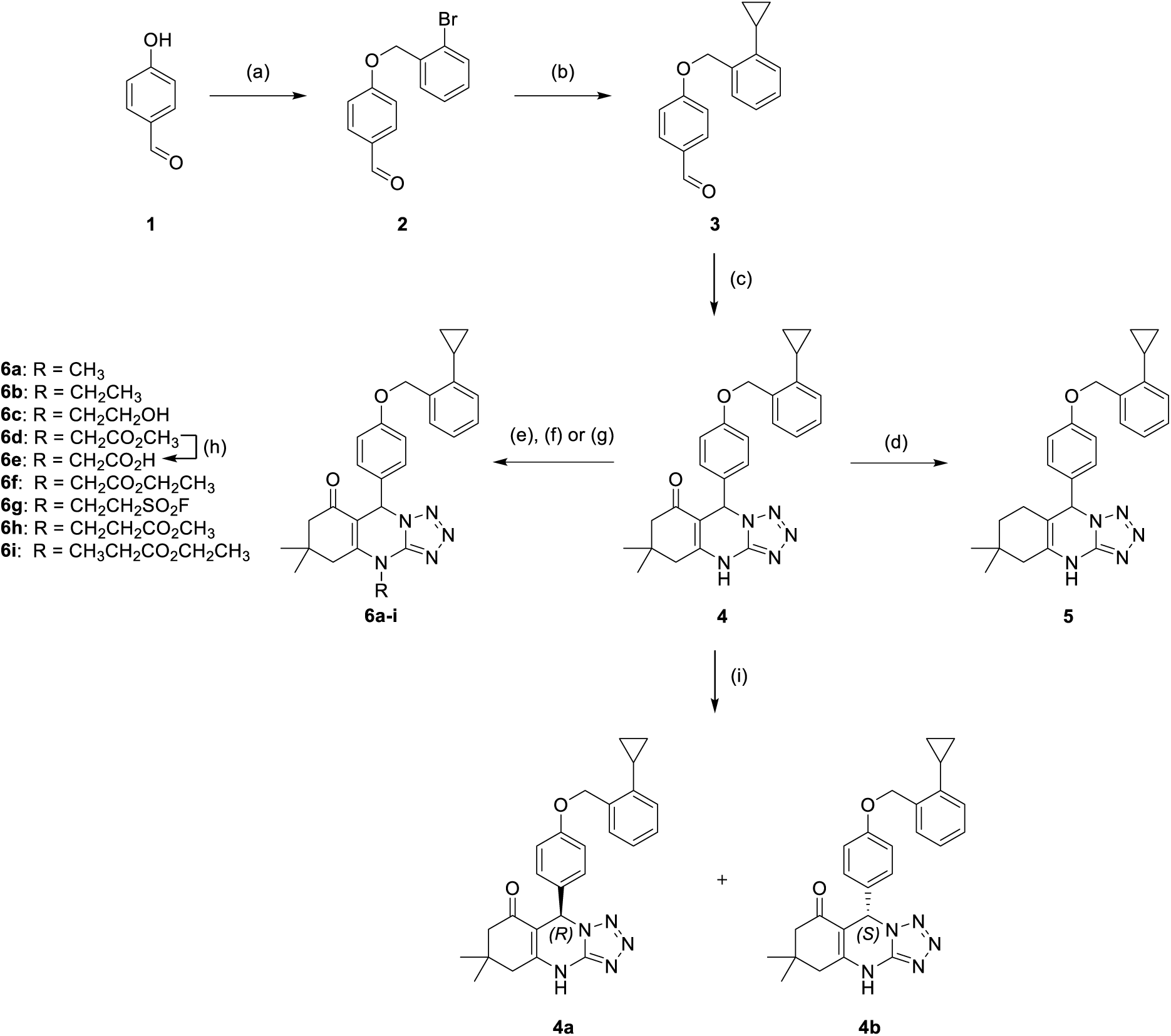
Synthetic pathway forming the tetrazoloquinazolinone 4 and its isolated enantiomers 4a,b, the deoxygenated species 5*, a*nd its *N*-substituted derivatives 6a-i*^a^*. ***^a^***Reagents and conditions: (a) 2-bromobenzyl bromide, DMF, Cs_2_CO_3_, rt, overnight, 94%; (b) cyclopropylboronic acid, Pd(OAc)_2_, P(C₆H₁₁)₃, K_3_PO_4_, toluene/H_2_O (95:5), 100 °C, 1 h, quantitative; (c) 5,5-dimethyl-1,3-cyclohexanedione, 5-aminotetrazole, MeOH, conc. H_2_SO_4_, reflux, overnight, 23%; (d) **4**, LiAlH_4_, dry THF, 0 °C – rt, overnight, 61% (**5**); (e) appropriate haloalkyl, Et_3_N, DMF (for **6a-c**), MeCN (for **6d**,**f**), rt – 60 °C, 3 – 4 d, 41% (**6a**), 73% (**6b**), 33% (**6c**), 96% (**6d**), 60% (**6f**); (f) ethenesulfonyl fluoride, dry DCM, rt – 50 °C, 4 d, 42% (**6g**); (g) appropriate Michael acceptor, Et_3_N, DMF, MeCN, reflux, 3 d, 71% (**6h**), 52% (**6i**); (h) i. LiOH.H_2_O, THF/H_2_O (1:1), ii. citric acid, rt, 3 h, quantitative; (i) **4**, Chiral HPLC, cellulose-2 packing, 65% (**4a**), 70% (**4b**).

#### Experimental

All reagents and solvents used were obtained from standard suppliers and used without additional purification. Reaction progress was monitored *via* thin-layer chromatography (TLC) utilizing Merck Millipore TLC silica gel 60 F_254_ aluminum plates, which were visualized under ultraviolet light (254 nm). Flash column chromatography (FCC) was performed using either isocratic or gradient solvent ratios through Merck silica gel 60, particle size 40-63 μm. ^1^H NMR, ^13^C NMR and ^19^F NMR spectra were recorded using a Bruker Avance III Nanobay 400 MHz spectrometer equipped with a BACS 60 automatic sample changer at 400 MHz, 101 MHz and 377 MHz, respectively. All chemical shifts (*δ*) are recorded in parts per million (ppm), with all peaks being referenced through the residual deuterated solvent peak. Experimental data was recorded in the following format: chemical shift (multiplicity, coupling constant, integration) whereby multiplicity is denoted by; singlet (s), doublet (d), triplet (t), quartet (q), pentet (p), heptet (hept), doublet of doublets (dd), triplet of doublets (td), doublet of doublets of doublets (ddd), and multiplet (m). Coupling constants were recorded as *J* in Hertz (Hz). ^13^C NMR assignments were based on the spectral phasing of carbon environments in regard to the APT test where, C = quaternary carbon, CH = methine carbon, CH_2_ = methylene carbon and CH_3_ = methyl carbon. Deuterated solvents were purchased from Cambridge Isotope Laboratories, Inc. LC-MS analysis was conducted on an Agilent 1260 LCMS SQ equipped with a 1260 Infinity G1312B Binary pump and a 1260 Infinity G1367E 1260 HiP ALS autosampler. Detection of UV reactive compounds was performed at wavelengths of 214 nm and 254 nm and were recorded by a 1290 Infinity G4212A 1290 DAD variable wavelength detector. LC-MS data was processed through the LC/MSD Chemstation Rev.B.04.03 SP2 coupled with MassHunter Easy Access Software. The LC component was run as a reverse phase HPLC using a Raptor C18 2.7 μm 50 × 3.0 mm column at 35 °C. Solvents comprised of buffer A was 0.1% formic acid in H_2_O, and buffer B was 0.1% formic acid in MeCN. The gradient was: 0−1 min 95% buffer A and 5% buffer B, from 1 to 2.5 min up to 0% buffer A and 100% buffer B, held at this composition until 3.8 min, 3.8−4 min 95% buffer A and 5% buffer B, held until 5 min at this composition; the flow rate was 0.5 mL/min and total run time was 5 min. The mass spectrum section was acquired in positive and negative ion mode using a scan range of 100 – 1000 *m/z*. HR-MS analysis was conducted with an Agilent 6224 TOF LC/MS Mass Spectrometer coupled to an Agilent 1290 Infinity (Agilent, Palo Alto, CA). All data were acquired and reference mass corrected via a dual-spray electrospray ionization (ESI) source. Each scan or data point on the Total Ion Chromatogram (TIC) is an average of 13,700 transients, producing a spectrum every second. Mass spectra were created by averaging the scans across each peak and background subtracted against the first 10 seconds of the TIC. Mass spectrometer conditions used were as follows; electrospray ionization mode, 11 L/min drying gas flow, 45 psi nebulizer, 325 °C drying gas temperature, 4000 V capillary voltage, 160 V fragmentor, 65 V skimmer, 750 V OCT RFV and a scan range of 100-1500 *m/z*. The internal reference ions were *m/z* = 121.050873 & 922.009798 in the positive ion mode. Analytical HPLC was performed on an Agilent 1260 analytical HPLC system through a Zorbax Eclipse Plus C18 Rapid Resolution 4.6 × 100 mm 3.5 Micron column using a flow rate of 20 mL/min and a gradient of 5 – 100% B over 12 min followed by 100% B over 3 min where: A = Milli-Q water and B = HPLC grade MeCN.

#### 4-((2-Bromobenzyl)oxy)benzaldehyde (2)

To a solution of *p*-hydroxybenzaldehyde (400 mg, 3.28 mmol, 1.0 equiv.) in DMF (16 mL) was added 2-bromobenzyl bromide (819 mg, 3.28 mmol, 1.0 equiv.) and Cs_2_CO_3_ (2.14 g, 6.56 mmol, 2.0 equiv.). The reaction mixture was stirred at room temperature for 20 h, after which LC-MS analysis was conducted to show conversion of starting material to a single product. The reaction mixture was partitioned between EtOAc (40 mL) and water (40 mL), and the phases were separated. The organic phase was further washed with water (2 × 40 mL) followed by sat. NaCl (40 mL), dried over MgSO_4_, gravity filtered then concentrated *in vacuo* to yield the desired product as a yellow oil (897 mg, 94%). LC-MS *(m/z):* 293.0 [M+H]^+^. TLC *R_f_* = 0.72 (PET/EtOAc, 1:1). ^1^H NMR (DMSO-*d*_6_) *δ* 9.89 (s, 1H), 7.93 – 7.87 (m, 2H), 7.70 (dd, *J* = 1.2, 8.0 Hz, 1H), 7.61 (dd, *J* = 1.7, 7.6 Hz, 1H), 7.44 (td, *J* = 1.3, 7.5 Hz, 1H), 7.34 (td, *J* = 1.8, 7.7 Hz, 1H), 7.26 – 7.21 (m, 2H), 5.25 (s, 2H). ^13^C NMR (DMSO-*d*_6_) *δ* 191.8 (CH), 163.5 (C), 135.6 (C), 133.2 (CH), 132.3 (CH), 131.0 (CH), 130.9 (CH), 130.4 (C), 128.4 (CH), 123.5 (C), 115.6 (CH), 70.0 (CH_2_).

#### 4-((2-Cyclopropylbenzyl)oxy)benzaldehyde (3)

To a solution of 4-((2-bromobenzyl)oxy)benzaldehyde (**2**) (128 mg, 440 µmol, 1.0 equiv.), cyclopropylboronic acid (49 mg, 571 µmol, 1.3 equiv.), K_3_PO_4_ (327 mg, 1.54 mmol, 3.5 equiv.) and tricyclohexylphosphine (12 mg, 44 mmol, 0.10 equiv.) in toluene (2.5 mL) and water (125 µL) under a nitrogen atmosphere was added Pd(OAc)_2_ (4.9 mg, 22 mmol, 0.05 equiv.). The mixture was heated to 100 °C for 1 h and then cooled to room temperature. The crude mixture was partitioned between water (40 mL) and EtOAc (40 mL), and the phases were separated. The organic phase was further washed with water (2 × 40 mL) followed by sat. NaCl (40 mL), dried over MgSO_4_ and gravity filtered then concentrated *in vacuo* to afford the crude product as a yellow oil (113 mg, quantitative). LC-MS *(m/z):* No observable mass trace. TLC *R_f_* = 0.75 (PET/EtOAc, 1:1). ^1^H NMR (CDCl_3_) *δ* 9.95 (s, 1H), 7.96 – 7.86 (m, 2H), 7.46 (dd, *J* = 1.5, 7.5 Hz, 1H), 7.34 (td, *J* = 1.6, 7.5 Hz, 1H), 7.28 (td, *J* = 1.6, 7.5 Hz, 1H), 7.19 – 7.13 (m, 3H), 5.38 (s, 2H), 2.03 (tt, *J* = 5.4, 8.3 Hz, 1H), 1.08 – 0.96 (m, 2H), 0.82 – 0.71 (m, 2H). ^13^C NMR (CDCl_3_) *δ* 190.9 (CH), 164.1 (C), 141.5 (C), 135.2 (C), 132.1 (CH), 130.2 (C), 128.7 (CH), 128.4 (CH), 126.3 (CH), 126.2 (CH), 115.2 (CH), 68.6 (CH_2_), 12.6 (CH), 7.1 (CH_2_).

#### 9-(4-((2-Cyclopropylbenzyl)oxy)phenyl)-6,6-dimethyl-5,6,7,9-tetrahydrotetrazolo[5,1-*b*]quinazolin-8(4*H*)-one (4)

To a solution of 5,5-dimethyl-1,3-cyclohexanedione (47 mg, 339 µmol, 1.0 equiv.) and 1*H*-tetrazol-5-amine (29 mg, 339 µmol, 1.0 equiv.) in MeOH (1 mL) was added 4-((2-cyclopropylbenzyl)oxy)benzaldehyde (**3**) (85 mg, 339 µmol, 1.0 equiv.) and 98% H_2_SO_4_ (10 µL). The reaction mixture was left to stir under reflux overnight, after which TLC analysis (PET/EtOAc, 1:1) showed total consumption of starting material. The precipitate which formed was collected by suction filtration and washed with cold MeOH (∼20 mL) to afford the crude product which was purified *via* preparative reverse-phase HPLC to afford the desired product as a white powder (34 mg, 23%). LC-MS *(m/z):* 442.2 [M+H]^+^. TLC *R_f_* = 0.52 (PET/EtOAc, 1:1). HPLC: *t*_R_ 5.74 min, >95% purity (254 nm). HRMS: calcd for C_26_H_28_N_5_O_2+_ [M+H]^+^ 442.2238, found 442.2250. ^1^H NMR (DMSO-*d*_6_) *δ* 11.57 (s, 1H), 7.37 (dd, *J* = 1.5, 7.5 Hz, 1H), 7.29 – 7.20 (m, 3H), 7.17 (td, *J* = 1.4, 7.5 Hz, 1H), 7.04 – 6.92 (m, 3H), 6.56 (s, 1H), 5.19 (s, 2H), 2.59 (s, 2H), 2.23 (d, *J* = 16.2 Hz, 1H), 2.13 (d, *J* = 16.2 Hz, 1H), 2.00 (tt, *J* = 5.3, 8.4 Hz, 1H), 1.06 (s, 3H), 1.01 (s, 3H), 0.94 – 0.84 (m, 2H), 0.70 – 0.57 (m, 2H). ^13^C NMR (DMSO-*d*_6_) *δ* 193.4 (C), 158.8 (C), 150.9 (C), 148.9 (C), 142.0 (C), 136.0 (C), 133.4 (C), 128.9 (CH), 128.9 (CH), 128.7 (CH), 115.1 (CH), 106.1 (C), 68.1 (CH_2_), 57.4 (CH), 50.3 (CH_2_), 32.7 (C), 28.7 (CH_3_), 27.5 (CH_3_), 12.3 (CH_3_), 7.8 (CH_2_).

#### 9-(4-((2-Cyclopropylbenzyl)oxy)phenyl)-6,6-dimethyl-4,5,6,7,8,9-hexahydrotetrazolo[5,1-*b*]quinazoline (5)

To an oven dried reaction vessel under an N_2_ atmosphere was added 9-(4-((2-cyclopropylbenzyl)oxy)phenyl)-6,6-dimethyl-5,6,7,9-tetrahydrotetrazolo[5,1-*b*]quinazolin-8(4*H*)-one (**4**) (250 mg, 566 μmol, 1.0 equiv.) in dry THF (10 mL) followed by the dropwise addition of LiAlH_4_ (1 M in THF) (1.13 mL, 1.13 mmol, 2.0 equiv.) at 0 °C. The reaction mixture was left to stir at rt for 20 h after which it was quenched by the addition of Rochelle’s salt (20 mL, 10% w/w). The reaction mixture was partitioned between EtOAc (50 mL) and water (50 mL), and the phases were separated. The organic phase was further washed with water (2 × 40 mL) followed by brine (50 mL), dried over MgSO_4_, gravity filtered then concentrated *in vacuo*. The residue was purified *via* FCC (PET/EtOAc 3:1), whereby the desired eluates were reduced *in vacuo* to afford the desired product as a white solid (147 mg, 61%). LC-MS *(m/z):* 428.2 [M+H]^+^. TLC *R_f_* = 0.28 (PET/EtOAc, 3:1). HPLC: *t_R_* 8.82 min, >95% purity (254 nm). HRMS: calcd for C_26_H_29_N_5_ONa^+^ [M+Na]^+^ 428.2445, found 428.2454. ^1^H NMR (DMSO-*d*_6_) *δ* 9.91 (s, 1H), 7.39 (dd, *J* = 1.5, 7.5 Hz, 1H), 7.25 (td, *J* = 1.6, 7.5 Hz, 1H), 7.23 – 7.13 (m, 3H), 7.09 – 7.01 (m, 2H), 7.05 – 6.97 (m, 1H), 6.12 (s, 1H), 5.22 (s, 2H), 2.10 – 1.95 (m, 3H), 1.87 (dt, *J* = 5.3, 16.2 Hz, 1H), 1.62 – 1.48 (m, 1H), 1.43 – 1.26 (m, 2H), 0.96 (s, 3H), 0.93 – 0.87 (m, 2H), 0.86 (s, 3H), 0.69 – 0.62 (m, 2H). ^13^C NMR (DMSO-*d*_6_) *δ* 158.6 (C), 149.5 (C), 141.5 (C), 135.5 (C), 131.9 (C), 128.4 (CH), 128.4 (CH), 128.2 (CH), 127.2 (CH), 125.5 (CH), 124.9 (CH), 114.9 (CH), 100.8 (C), 67.7 (CH_2_), 61.2 (CH), 39.1 (CH_2_), 34.4 (CH_2_), 29.1 (CH_2_), 28.7 (CH_3_), 26.4 (CH_3_), 22.5 (C), 11.9 (CH), 7.4 (CH_2_).

#### 9-(4-((2-Cyclopropylbenzyl)oxy)phenyl)-4,6,6-trimethyl-5,6,7,9-tetrahydrotetrazolo[5,1-*b*]quinazolin-8(4*H*)-one (6a)

9-(4-((2-Cyclopropylbenzyl)oxy)phenyl)-6,6-dimethyl-5,6,7,9-tetrahydrotetrazolo[5,1-*b*]quinazolin-8(4*H*)-one (**4**) (80 mg, 181 μmol, 1.0 equiv.), iodomethane (113 μL, 1.81 mmol, 10 equiv.) and Et_3_N (253 μL, 1.81 mmol, 10 equiv.) were dissolved in DMF (3 mL) and stirred at rt overnight. A further 5.0 equiv. of iodomethane and 5.0 equiv. of Et_3_N were added and the reaction mixture was left to stir at rt for an additional 3 d. The reaction mixture was partitioned between EtOAc (50 mL) and water (40 mL), and the phases were separated. The organic phase was further washed with water (2 × 40 mL) followed by brine (40 mL), dried over MgSO_4_, gravity filtered then concentrated *in vacuo*. The residue was purified *via* preparative reverse-phase HPLC, resulting in a light-yellow solid (34 mg, 41%). LC−MS (*m*/*z):* 456.2 [M+H]^+^. TLC *R_f_* = 0.30 (PET/EtOAc, 1:1). HPLC: *t*_R_ 9.53 min, >95% purity (254 nm). HRMS: calcd for C_27_H_30_N_5_O_2+_ [M+H]^+^ 456.2394, found 456.2394. ^1^H NMR (DMSO-*d*_6_) *δ* 7.36 (dd, *J* = 1.5, 7.5 Hz, 1H), 7.28 – 7.21 (m, 3H), 7.16 (td, *J* = 1.4, 7.5 Hz, 1H), 7.00 (dd, *J* = 1.4, 7.9 Hz, 1H), 7.01 – 6.93 (m, 2H), 6.58 (s, 1H), 5.19 (s, 2H), 3.59 (s, 3H), 2.85 (d, *J* = 17.6 Hz, 1H), 2.70 (d, *J* = 17.6 Hz, 1H), 2.23 (d, *J* = 16.0 Hz, 1H), 2.12 (d, *J* = 16.0 Hz, 1H), 2.04 – 1.94 (m, 1H), 1.09 (s, 3H), 1.01 (s, 3H), 0.93 – 0.84 (m, 2H), 0.68 – 0.60 (m, 2H). ^13^C NMR (DMSO-*d*_6_) *δ* 193.1 (C), 158.4 (C), 151.7 (C), 150.5 (C), 141.5 (C), 135.5 (C), 132.8 (C), 128.5 (CH), 128.4 (CH), 128.2 (CH), 125.5 (CH), 124.8 (CH), 114.6 (CH), 107.1 (C), 67.7 (CH_2_), 56.3 (CH), 49.1 (CH_2_), 37.8 (CH_2_), 33.2 (CH_3_), 31.8 (C), 28.7 (CH_3_), 26.9 (CH_3_), 11.9 (CH), 7.4 (CH_2_).

#### 9-(4-((2-Cyclopropylbenzyl)oxy)phenyl)-4-ethyl-6,6-dimethyl-5,6,7,9-tetrahydrotetrazolo[5,1-*b*]quinazolin-8(4*H*)-one (6b)

9-(4-((2-Cyclopropylbenzyl)oxy)phenyl)-6,6-dimethyl-5,6,7,9-tetrahydrotetrazolo[5,1-*b*]quinazolin-8(4*H*)-one (**4**) (80 mg, 181 μmol, 1.0 equiv.), iodoethane (147 μL, 1.81 mmol, 10 equiv.) and Et_3_N (253 μL, 1.81 mmol, 10 equiv.) were dissolved in DMF (3 mL) and stirred at rt overnight. A further 5.0 equiv. of 2-bromoethan-1-ol and 5.0 equiv. of Et_3_N were added and the reaction mixture was left to stir at rt for an additional 3 d. The reaction mixture was partitioned between EtOAc (50 mL) and water (40 mL), and the phases were separated. The organic phase was further washed with water (2 × 40 mL) followed by brine (40 mL), dried over MgSO_4_, gravity filtered then concentrated *in vacuo*. The residue was purified *via* flash column chromatography (FCC) (PET/EtOAc 2:1 → 1:1), whereby the desired eluates were reduced *in vacuo* to afford the desired product as a white solid (62 mg, 73%). LC-MS *(m/z):* 470.2 [M+H]_+_. TLC *R_f_* = 0.20 (PET/EtOAc, 2:1). HPLC: *t_R_* 8.02 min, >95% purity (254 nm). HRMS: calcd for C_28_H_32_N_5_O_2+_ [M+H]^+^ 470.2515, found 470.2511. ^1^H NMR (CDCl_3_) *δ* 7.36 (dd, *J* = 1.5 Hz, 7.5, 1H), 7.24 (dd, *J* = 1.7, 7.5 Hz, 1H), 7.21 – 7.15 (m, 3H), 7.05 (dt, *J* = 1.0, 7.5 Hz, 1H), 6.94 – 6.88 (m, 2H), 6.68 (s, 1H), 5.17 (s, 2H), 4.26 – 4.06 (m, 2H), 2.68 (d, *J* = 17.1 Hz, 1H), 2.60 (d, *J* = 17.0 Hz, 1H), 2.31 (d, *J* = 16.3 Hz, 1H), 2.27 (d, *J* = 16.5 Hz, 1H), 1.99 – 1.89 (m, 1H), 1.44 (t, *J* = 7.1 Hz, 3H), 1.18 (s, 3H), 1.11 (s, 3H), 0.96 – 0.89 (m, 2H), 0.71 – 0.65 (m, 2H). ^13^C NMR (CDCl_3_) *δ* 193.5 (C), 159.2 (C), 150.0 (C), 149.3 (C), 141.4 (C), 135.7 (C), 132.1 (C), 128.3 (CH), 128.3 (CH), 128.2 (CH), 125.9 (CH), 125.8 (CH), 115.0 (CH), 109.2 (C), 68.2, 56.8 (CH), 49.5 (CH3), 41.7 (CH_3_), 38.6 (CH_3_), 32.5 (C), 29.2 (CH_3_), 27.6 (CH_3_), 14.4 (CH_3_), 12.3 (CH), 7.0 (CH_2_).

#### 9-(4-((2-Cyclopropylbenzyl)oxy)phenyl)-4-(2-hydroxyethyl)-6,6-dimethyl-5,6,7,9- tetrahydrotetrazolo[5,1-*b*]quinazolin-8(4*H*)-one (6c)

9-(4-((2-Cyclopropylbenzyl)oxy)phenyl)-6,6-dimethyl-5,6,7,9-tetrahydrotetrazolo[5,1-*b*]quinazolin-8(4*H*)-one (**4**) (50 mg, 113 μmol, 1.0 equiv.), 2-bromoethan-1-ol (8.0 μL, 113 μmol, 1.0 equiv.) and Et_3_N (15.8 μL, 113 μmol, 1.0 equiv.) were dissolved in DMF (2 mL) and stirred at rt overnight. A further 10 equiv. of 2-bromoethan-1-ol and 10 equiv. of Et_3_N were added and the reaction mixture was left to stir at rt for an additional 2 d. The reaction mixture was partitioned between EtOAc (30 mL) and water (30 mL), and the phases were separated. The organic phase was further washed with water (2 × 30 mL) followed by brine (30 mL), dried over MgSO_4_, gravity filtered then concentrated *in vacuo*. The residue was purified *via* FCC (PET/EtOAc 2:1), whereby the desired eluates were reduced *in vacuo* to afford the desired product as a white solid (18 mg, 33%). LC-MS *(m/z):* 486.1 [M+H]^+^. TLC *R_f_* = 0.31 (PET/EtOAc, 2:1). HPLC: *t_R_* 7.51 min, >95% purity (254 nm). HRMS: calcd for C_28_H_32_N_5_O_3+_ [M+H]^+^ 486.1884, found 486.1878. 1H NMR (CDCl_3_) *δ* 7.36 (dd, *J* = 1.5, 7.4 Hz, 1H), 7.23 (dd, *J* = 2.6, 9.1 Hz, 3H), 7.18 (td, *J* = 1.4, 7.4 Hz, 1H), 7.07 – 7.03 (m, 1H), 6.93 – 6.88 (m, 2H), 6.68 (s, 1H), 5.17 (s, 2H), 4.32 (dt, *J* = 4.8, 15.1 Hz, 1H), 4.20 (ddd, *J* = 4.2, 6.7, 15.1 Hz, 1H), 4.08 – 3.97 (m, 2H), 2.85 (d, *J* = 17.0 Hz, 1H), 2.67 (d, *J* = 17.0 Hz, 1H), 2.30 (d, *J* = 16.5 Hz, 1H), 2.25 (d, *J* = 16.6 Hz, 1H), 1.99 – 1.90 (m, 1H), 1.16 (s, 3H), 1.09 (s, 3H), 0.96 – 0.88 (m, 2H), 0.71 – 0.65 (m, 2H). ^13^C NMR (CDCl_3_) *δ* 194.1 (C), 159.3 (C), 150.6 (C), 150.3 (C), 141.5 (C), 135.9 (C), 132.2 (C), 128.5 (CH), 128.5 (CH), 128.4 (CH), 126.0 (CH), 125.9 (CH), 115.1 (CH), 109.1 (C), 68.4 (CH2), 60.4 (CH_2_), 57.0 (CH), 49.7 (CH_2_), 47.8 (CH_2_), 39.3 (CH_2_), 32.6 (C), 29.3 (CH_3_), 27.7 (CH_3_), 12.5 (CH), 7.2 (CH_2_).

#### Methyl-2-(9-(4-((2-cyclopropylbenzyl)oxy)phenyl)-6,6-dimethyl-8-oxo-5,7,8,9-tetrahydrotetrazolo[5,1-*b*]quinazolin-4(6*H*)-yl)acetate (6d)

9-(4-((2-Cyclopropylbenzyl)oxy)phenyl)-6,6-dimethyl-5,6,7,9-tetrahydrotetrazolo[5,1-*b*]quinazolin-8(4*H*)-one (**4**) (70 mg, 159 μmol, 1.0 equiv.), ethyl chloroacetate (20.8 μL, 238 μmol, 1.5 equiv.) and Et_3_N (33.2 μL, 238 μmol, 1.5 equiv.) were dissolved in MeCN (3 mL) and stirred at 60 °C overnight. A further 2.0 equiv. of ethyl chloroacetate and 2.0 equiv. of Et_3_N were added and the reaction mixture was left to stir at rt for an additional 3 d. The reaction mixture was partitioned between EtOAc (50 mL) and water (50 mL), and the phases were separated. The organic phase was further washed with water (3 × 40 mL) followed by brine (50 mL), dried over MgSO_4_, gravity filtered then concentrated *in vacuo* to afford the desired product as an off-white solid (78 mg, 96%). LC-MS *(m/z):* 514.2 [M+H]^+^. TLC *R_f_* = 0.62 (PET/EtOAc, 1:1). HPLC: *t_R_* 7.87 min, >95% purity (254 nm). HRMS: calcd for C_29_H_32_N_5_O_4+_ [M+H]^+^ 514.2422, found 514.2415. ^1^H NMR (CDCl_3_) *δ* 7.37 (dd, *J* = 1.6, 7.5 Hz, 1H), 7.34 – 7.30 (m, 2H), 7.24 (dd, *J* = 1.7, 7.5 Hz, 1H), 7.18 (td, *J* = 1.5, 7.4 Hz, 1H), 7.08 – 7.03 (m, 1H), 6.97 – 6.91 (m, 2H), 6.68 (s, 1H), 5.17 (s, 2H), 5.00 (d, *J* = 18.3 Hz, 1H), 4.76 (d, *J* = 18.3 Hz, 1H), 3.86 (s, 3H), 2.48 (s, 2H), 2.31 (d, *J* = 16.3 Hz, 1H), 2.27 (d, *J* = 16.5 Hz, 1H), 1.99 – 1.90 (m, 1H), 1.16 (s, 3H), 1.08 (s, 3H), 0.97 – 0.88 (m, 2H), 0.73 – 0.61 (m, 2H). ^13^C NMR (CDCl_3_) *δ* 193.8 (C), 168.2 (C), 159.4 (C), 150.4 (C), 148.8 (C), 141.5 (C), 135.9 (C), 131.8 (C), 128.6 (CH), 128.4 (CH), 128.4 (CH), 126.0 (CH), 125.9 (CH), 115.1 (CH), 110.2 (C), 68.3 (CH_2_), 57.2(CH), 53.3 (CH_3_), 49.7 (CH_2_), 46.9 (CH_2_), 38.7 (CH_2_), 32.6 (C), 29.3 (CH_3_), 27.6 (CH_3_), 12.5 (CH), 7.2 (CH_2_).

#### 2-(9-(4-((2-Cyclopropylbenzyl)oxy)phenyl)-6,6-dimethyl-8-oxo-5,7,8,9-tetrahydrotetrazolo[5,1-*b*]quinazolin-4(6*H*)-yl)acetic acid (6e)

Methyl 2-(9-(4-((2-cyclopropylbenzyl)oxy)phenyl)-6,6-dimethyl-8-oxo-5,7,8,9-tetrahydrotetrazolo[5,1-*b*]quinazolin-4(6*H*)-yl)acetate (**6d**) (120 mg, 234 µmol, 1.0 equiv.) and LiOH.H_2_O (98 mg, 2.34 mmol, 10.0 equiv.) was stirred at rt in a THF/H_2_O mixture (1:1, 4mL) for 3 h. The reaction mixture was concentrated *in vacuo* and partitioned between EtOAc (40 mL) and 1 M citric acid (40 mL), and the phases were separated. The organic phase was further washed with water (3 × 40 mL) followed by brine (40 mL), dried over MgSO_4_, gravity filtered then concentrated *in vacuo* to afford the desired product as a white solid (116 mg, quant.). LC-MS *(m/z):* 500.2 [M+H]^+^. TLC *R_f_* = 0.31 (PET/EtOAc, 2:1). HPLC: *t_R_* 7.51 min, >95% purity (254 nm). HRMS: calcd. for C_28_H_29_N_5_O_3+_ [M+H]^+^ 500.2292, found 500.2295. ^1^H NMR (DMSO-*d*_6_) *δ* 12.30 (s, 1H), 7.36 (dd, *J* = 1.5, 7.5 Hz, 1H), 7.36 – 7.27 (m, 2H), 7.24 (td, *J* = 1.6, 7.6 Hz, 1H), 7.16 (td, *J* = 1.4, 7.4 Hz, 1H), 7.06 – 6.93 (m, 3H), 6.63 (s, 1H), 5.19 (s, 2H), 4.90 (d, *J* = 18.5 Hz, 1H), 4.85 (d, *J* = 18.5 Hz, 1H), 2.76 (d, *J* = 15.4 Hz, 1H), 2.65 (d, *J* = 15.4 Hz, 1H), 2.27 (d, *J* = 16.2 Hz, 1H), 2.15 (d, *J* = 16.1 Hz, 1H), 2.04 – 1.93 (m, 1H), 1.07 (s, 3H), 0.98 (s, 3H), 0.92 – 0.85 (m, 2H), 0.69 – 0.60 (m, 2H). ^13^C NMR (DMSO-*d*_6_) *δ* 193.6 (C), 169.4 (C), 158.6 (C), 151.2 (C), 150.4 (C), 141.7 (C), 135.6 (C), 132.9 (C), 128.7 (CH), 128.6 (CH), 128.4 (CH), 125.6 (CH), 125.0 (CH), 114.7 (CH), 108.0 (C), 67.8 (CH_2_), 56.5 (CH), 49.2 (CH_2_), 47.8 (CH_2_), 37.5 (CH_2_), 32.1 (C), 28.7 (CH_3_), 27.0 (CH_3_), 12.0 (CH), 7.5 (CH_2_).

#### Ethyl-2-(9-(4-((2-cyclopropylbenzyl)oxy)phenyl)-6,6-dimethyl-8-oxo-5,7,8,9-tetrahydrotetrazolo[5,1-*b*]quinazolin-4(6*H*)-yl)acetate (6f)

9-(4-((2-Cyclopropylbenzyl)oxy)phenyl)-6,6-dimethyl-5,6,7,9-tetrahydrotetrazolo[5,1-*b*]quinazolin-8(4*H*)-one (**4**) (100 mg, 226 μmol, 1.0 equiv.), ethyl chloroacetate (72.7 μL, 679 μmol, 3.0 equiv.) and Et_3_N (94.7 μL, 679 μmol, 3.0 equiv.) were dissolved in MeCN (4 mL) and stirred at 60 °C overnight. A further 1.0 equiv. of ethyl chloroacetate was added and the reaction mixture was left to stir at rt for an additional 3 d. The reaction mixture was partitioned between EtOAc (50 mL) and water (50 mL), and the phases were separated. The organic phase was further washed with water (3 × 40 mL) followed by brine (50 mL), dried over MgSO_4_, gravity filtered then concentrated *in vacuo*. The residue was purified *via* FCC (PET/EtOAc 2:1 → 1:1), whereby the desired eluates were reduced *in vacuo* to afford the desired product as a white solid (71 mg, 60%). LC-MS *(m/z):* 528.2 [M+H]^+^. TLC *R_f_* = 0.25 (PET/EtOAc, 2:1). HPLC: *t_R_* 8.39 min, >95% purity (254 nm). HRMS: calcd for C_30_H_34_N_5_O_4+_ [M+H]^+^ 528.2605, found 528.2619. ^1^H NMR (CDCl_3_) *δ* 7.37 (dd, *J* = 1.6, 7.5 Hz, 1H), 7.34 – 7.30 (m, 2H), 7.24 (dd, *J* = 1.6, 7.5 Hz, 1H), 7.18 (td, *J* = 1.4, 7.4 Hz, 1H), 7.07 – 7.03 (m, 1H), 6.96 – 6.91 (m, 2H), 6.68 (s, 1H), 5.17 (s, 2H), 5.00 (d, *J* = 18.3 Hz, 1H), 4.73 (d, *J* = 18.3 Hz, 1H), 4.32 (q, *J* = 7.1 Hz, 2H), 2.48 (br s, 2H), 2.31 (d, *J* = 16.4 Hz, 1H), 2.26 (d, *J* = 16.3 Hz, 1H), 1.99 – 1.90 (m, 1H), 1.34 (t, *J* = 7.1 Hz, 3H), 1.16 (s, 3H), 1.07 (s, 3H), 0.97 – 0.88 (m, 2H), 0.70 – 0.65 (m, 2H). ^13^C NMR (CDCl_3_) *δ* 193.6 (C), 167.5 (C), 159.3 (C), 150.2 (C), 148.8 (C), 141.4 (C), 135.8 (C), 131.8 (C), 128.5 (CH), 128.3 (CH), 128.3 (CH), 125.9 (CH), 125.8 (CH), 115.0 (CH), 110.0 (C), 68.2 (CH_2_), 62.6 (CH_2_), 57.0 (CH), 49.5 (CH_2_), 46.9 (CH_2_), 38.6 (CH_2_), 32.51 (CH_2_), 29.2 (CH_3_), 27.4 (CH_3_), 14.1 (CH_3_), 12.3 (CH), 7.0 (CH_2_).

#### 2-(9-(4-((2-Cyclopropylbenzyl)oxy)phenyl)-6,6-dimethyl-8-oxo-5,7,8,9-tetrahydrotetrazolo[5,1-*b*]quinazolin-4(6*H*)-yl)ethane-1-sulfonyl fluoride (6g)

To a solution of 9-(4-((2-cyclopropylbenzyl)oxy)phenyl)-6,6-dimethyl-5,6,7,9-tetrahydrotetrazolo[5,1-*b*]quinazolin-8(4*H*)-one (**4**) (70 mg, 159 μmol, 1.0 equiv.) in dry DCM (1 mL) was added ethenesulfonyl fluoride (13.1 μL, 159 μmol, 1.0 equiv.). The reaction mixture was left to stir at rt overnight after which a further 2.0 equiv. of ethenesulfonyl fluoride was added and the temperature increased to 50 °C. The reaction was stopped after a further 3 d and subsequently concentrated *in vacuo*. The crude residue was purified *via* FCC (PET/EtOAc 9:1 → 2:1 → 1:1), whereby the desired eluates were reduced *in vacuo* to afford the desired product as a white solid (37 mg, 42%) following concentration. LC-MS *(m/z):* 552.2 [M+H]^+^. TLC *R_f_* = 0.24 (PET/EtOAc, 2:1). HPLC: *t_R_* 8.35 min, >95% purity (254 nm). HRMS: calcd for C_28_H_31_FN_5_O_4_S^+^ [M+H]^+^ 552.2442, found 552.2438. ^1^H NMR (CDCl_3_) *δ* 7.37 (dd, *J* = 1.6, 7.5 Hz, 1H), 7.27 – 7.22 (m, 3H), 7.19 (td, *J* = 1.5, 7.4 Hz, 1H), 7.05 (dd, *J* = 1.8, 7.5 Hz, 1H), 6.97 – 6.91 (m, 2H), 6.68 (d, *J* = 1.3 Hz, 1H), 5.17 (s, 2H), 4.66 (ddd, *J* = 3.9, 9.6, 15.6 Hz, 1H), 4.60 – 4.50 (m, 1H), 4.31 (dddd, *J* = 2.7, 4.6, 9.5, 14.3 Hz, 1H), 3.86 (dq, *J* = 4.3, 15.4 Hz, 1H), 2.85 (d, *J* = 17.0 Hz, 1H), 2.59 (d, *J* = 17.0 Hz, 1H), 2.30 (br s, 2H), 1.99 – 1.90 (m, 1H), 1.20 (s, 3H), 1.12 (s, 3H), 0.95 – 0.86 (m, 2H), 0.71 – 0.65 (m, 2H). ^13^C NMR (CDCl_3_) *δ* 193.8 (C), 159.5 (C), 149.1 (C), 148.4 (C), 141.5 (C), 135.8 (C), 131.3 (C), 128.6 (CH), 128.4 (CH), 128.4 (CH), 126.0 (CH), 125.9 (CH), 115.2 (CH), 110.3 (C), 68.3 (CH_2_), 57.3 (CH), 49.5 (CH_2_), 48.4 (d, *J* = 16.6 Hz, CH_2_), 40.2 (CH_2_), 38.6 (CH_2_), 32.5 (C), 29.8 (CH_3_), 27.1 (CH_3_), 12.5 (CH), 7.2 (CH_2_).

#### Methyl 3-(9-(4-((2-cyclopropylbenzyl)oxy)phenyl)-6,6-dimethyl-8-oxo-5,7,8,9-tetrahydrotetrazolo[5,1-*b*]quinazolin-4(6*H*)-yl)propanoate (6h)

9-(4-((2-Cyclopropylbenzyl)oxy)phenyl)-6,6-dimethyl-5,6,7,9-tetrahydrotetrazolo[5,1-*b*]quinazolin-8(4*H*)-one (**4**) (80 mg, 181 μmol, 1.0 equiv.), methyl acrylate (163 μL, 1.81 mmol, 10 equiv.) and Et_3_N (253 μL, 1.81 mmol, 10 equiv.) were dissolved in DMF (3 mL) and stirred at rt overnight. A further 5.0 equiv. of methyl acrylate and 5.0 equiv. of Et_3_N were added and the reaction mixture was left to stir at rt for an additional 3 d. The reaction mixture was partitioned between EtOAc (50 mL) and water (40 mL), and the phases were separated. The organic phase was further washed with water (2 × 40 mL) followed by brine (40 mL), dried over MgSO_4_, gravity filtered then concentrated *in vacuo*. The desired product was isolated *via* FCC (PET/EtOAc, 2:1 → 1:1), resulting in a white solid (68 mg, 71%). LC-MS *(m/z):* 528.3 [M+H]^+^. TLC *R_f_* = 0.25 (PET/EtOAc, 2:1). HPLC: *t_R_* 7.76 min, >95% purity (254 nm). HRMS: calcd. for C_30_H_34_N_5_O_4+_ [M+H]^+^ 528.2605, found 528.2607. ^1^H NMR (DMSO-*d*_6_) *δ* 7.36 (dd, *J* = 1.5, 7.5 Hz, 1H), 7.29 – 7.21 (m, 1H), 7.23 – 7.20 (m, 2H), 7.16 (td, *J* = 1.4, 7.4 Hz, 1H), 7.04 – 6.96 (m, 1H), 7.01 – 6.93 (m, 2H), 6.58 (s, 1H), 5.19 (s, 2H), 4.43 – 4.21 (m, 2H), 3.60 (s, 3H), 2.97 – 2.82 (m, 3H), 2.77 – 2.68 (m, 1H), 2.24 (d, *J* = 16.1 Hz, 1H), 2.14 (d, *J* = 16.1 Hz, 1H), 2.06 – 1.93 (m, 1H), 1.09 (s, 3H), 1.01 (s, 3H), 0.92 – 0.85 (m, 2H), 0.69 – 0.61 (m, 2H). ^13^C NMR (DMSO-*d*_6_) *δ* 193.3 (C), 171.0 (C), 158.5 (C), 150.8 (C), 149.6 (C), 141.5 (C), 135.5 (C), 132.6 (C), 128.4 (CH), 128.4 (CH), 128.2 (CH), 125.5 (CH), 124.8 (CH), 114.6 (CH), 107.8 (C), 67.7 (CH_2_), 56.2 (CH), 51.7 (CH_3_), 49.1 (CH_2_), 41.8 (CH_2_), 37.3 (CH_2_), 32.3 (CH_2_), 32.0 (C), 28.5 (CH_3_), 26.8 (CH_3_), 11.9 (CH), 7.4 (CH_2_).

#### Ethyl 3-(9-(4-((2-cyclopropylbenzyl)oxy)phenyl)-6,6-dimethyl-8-oxo-5,7,8,9-tetrahydrotetrazolo[5,1-*b*]quinazolin-4(6*H*)-yl)propanoate (5i)

9-(4-((2-Cyclopropylbenzyl)oxy)phenyl)-6,6-dimethyl-5,6,7,9-tetrahydrotetrazolo[5,1-*b*]quinazolin-8(4*H*)-one (**4**) (80 mg, 181 μmol, 1.0 equiv.), ethyl acrylate (193 μL, 1.81 mmol, 10 equiv.) and Et_3_N (253 μL, 1.81 mmol, 10 equiv.) were dissolved in DMF (3 mL) and stirred at rt overnight. A further 5.0 equiv. of ethyl acrylate and 5.0 equiv. of Et_3_N were added and the reaction mixture was left to stir at rt for an additional 3 d. The reaction mixture was partitioned between EtOAc (50 mL) and water (40 mL) and the phases were separated. The organic phase was further washed with water (2 × 40 mL) followed by brine (40 mL), dried over MgSO_4_, gravity filtered then concentrated *in vacuo*. The desired product was isolated *via* FCC (PET/EtOAc, 1:1), resulting in an off-white solid (51 mg, 52%). LC-MS *(m/z):* 542.3 [M+H]^+^. TLC *R_f_* = 0.22 (PET/EtOAc, 1:1). HPLC: *t_R_* 7.99 min, >95% purity (254 nm). HRMS: calcd. for C_31_H_36_N_5_O_4+_ [M+H]^+^ 542.2762, found 542.2761. ^1^H NMR (DMSO-*d*_6_) *δ* 7.36 (dd, *J* = 1.5, 7.5 Hz, 1H), 7.27 – 7.20 (m, 3H), 7.16 (td, *J* = 1.4, 7.4 Hz, 1H), 7.02 – 6.98 (m, 1H), 6.98 – 6.94 (m, 2H), 6.58 (s, 1H), 5.19 (s, 2H), 4.42 – 4.21 (m, 2H), 4.12 – 4.00 (m, 2H), 2.93 (d, *J* = 17.3 Hz, 1H), 2.85 (q, *J* = 7.3 Hz, 2H), 2.74 (d, *J* = 17.5 Hz, 1H), 2.24 (d, *J* = 16.1 Hz, 1H), 2.14 (d, *J* = 16.0 Hz, 1H), 2.05 – 1.94 (m, 1H), 1.14 (t, *J* = 7.1 Hz, 3H), 1.09 (s, 3H), 1.01 (s, 3H), 0.92 – 0.84 (m, 2H), 0.67 – 0.61 (m, 2H). ^13^C NMR (DMSO-*d*_6_) *δ* 194.4 (C), 171.2 (C), 158.9 (C), 151.7 (C), 150.1 (C), 142.0 (C), 135.8 (C), 132.9 (C), 129.0 (CH), 128.9 (CH), 128.8 (CH), 126.0 (CH), 125.4 (CH), 115.1 (CH), 108.3 (C), 68.2 (CH_2_), 61.0 (CH_2_), 56.7 (CH), 49.4 (CH_2_), 42.2 (CH_2_), 37.8 (CH_2_), 33.0 (CH_2_), 32.4 (C), 29.0 (CH_3_), 27.1 (CH_3_), 14.3 (CH_3_), 12.3 (CH), 7.8 (CH_2_).

---

**Supplementary Fig. 1.**
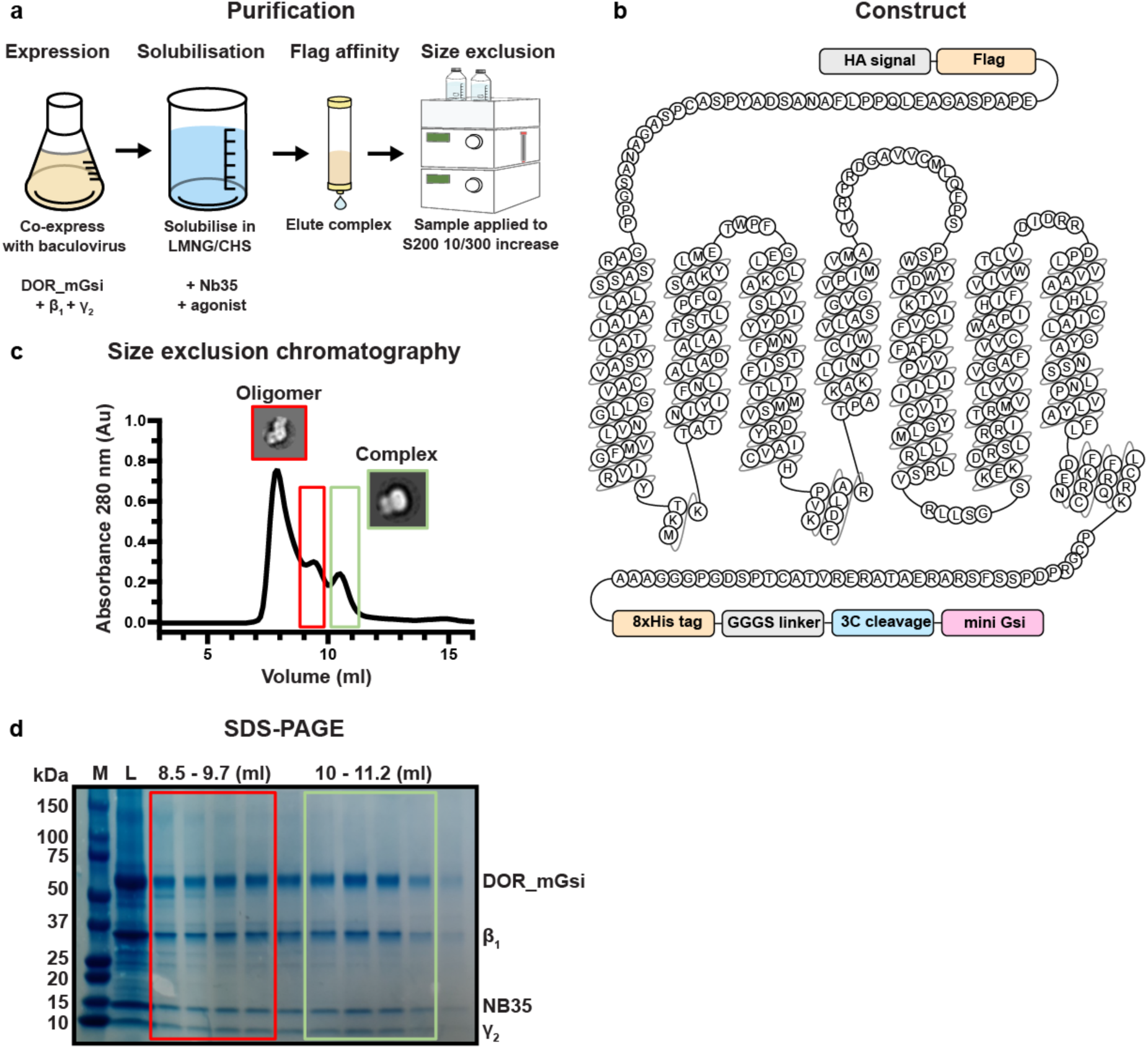
Purification and characterization of δOR-mG_si_-DADLE complex. (a) Schematic overview of the purification protocol used to isolate the δOR-mG_si_-DADLE complex. (b) Schematic representation of the δOR-mG_si_ fusion construct used for structural studies. (c) Size exclusion chromatography profile showing separation of oligomeric and monomeric forms of the δOR complex. (d) SDS-PAGE analysis of fractions across the size exclusion chromatography run. The gel shows molecular weight markers (M), load (L), and fractions from 8.5-9.7 ml (red box, oligomer fractions) and 10-11.2 mL (green box, monomeric complex fractions). Protein bands corresponding to δOR-mG_si_ (∼65 kDa), Gβ_1_(∼37 kDa), Nb35 (∼15 kDa), and Gγ_2_ (∼8 kDa) are indicated.

**Supplementary Fig. 2.**
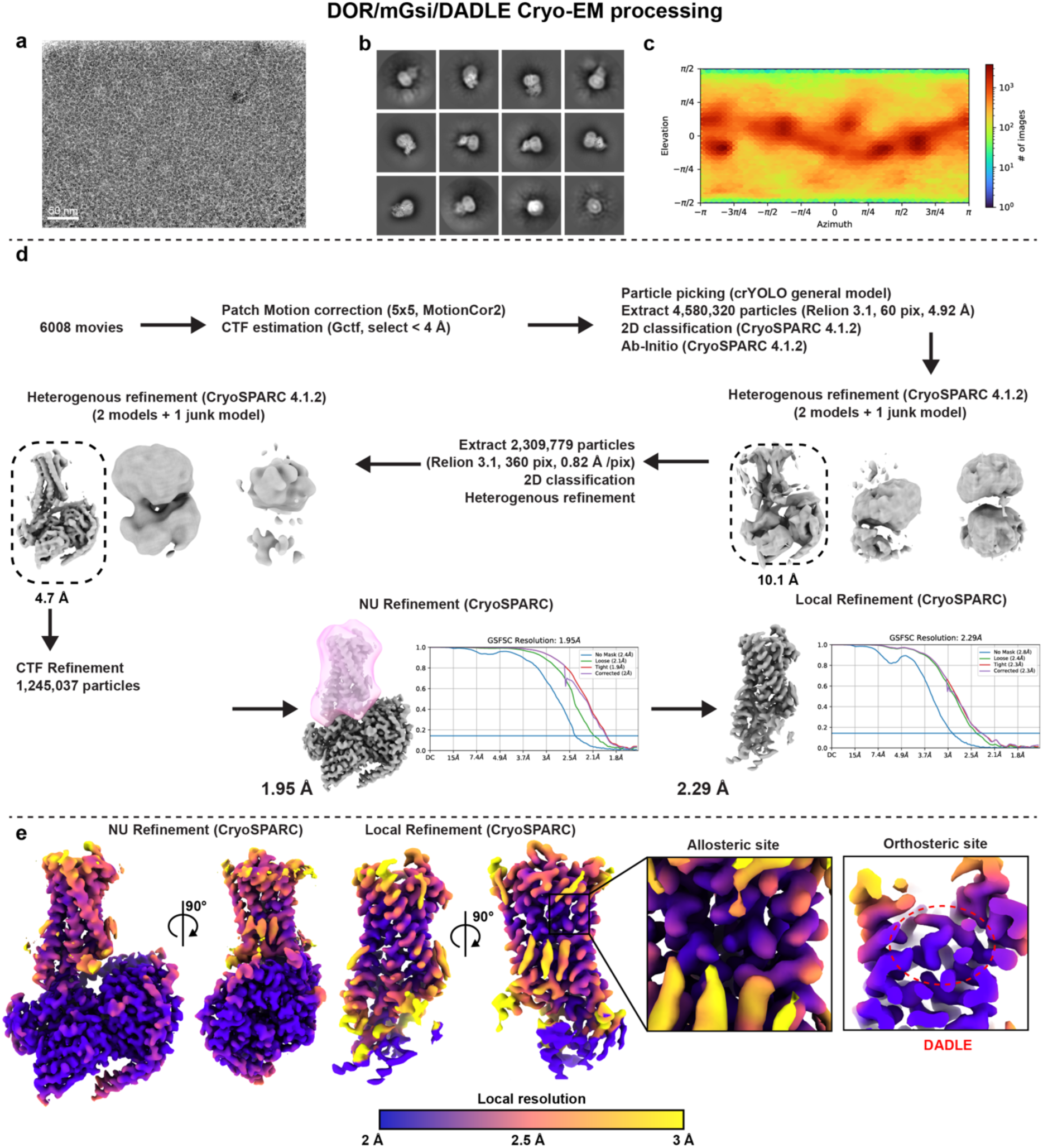
Cryo-EM data collection and processing workflow for δOR-mG_si_-DADLE complex. (a) Representative cryo-EM micrograph of the δOR-mG_si_-DADLE complex. Scale bar = 50 nm. (b) Gallery of 2D class averages from reference-free classification showing different orientations of the δOR-mG_si_-DADLE complex. (c) Angular distribution plot showing the coverage of particle orientations used in the final 3D reconstruction. (d) Comprehensive data processing workflow starting from 6,008 raw movies to a final 3D reconstruction with 1,245,037 particles. (e) Final 3D reconstruction colored by local resolution, showing two different views of the consensus (NU refinement) and receptor-focused (Local refinement) maps. Insets show detailed views of the allosteric binding site (left) and orthosteric binding site containing DADLE (right).

**Supplementary Fig. 3.**
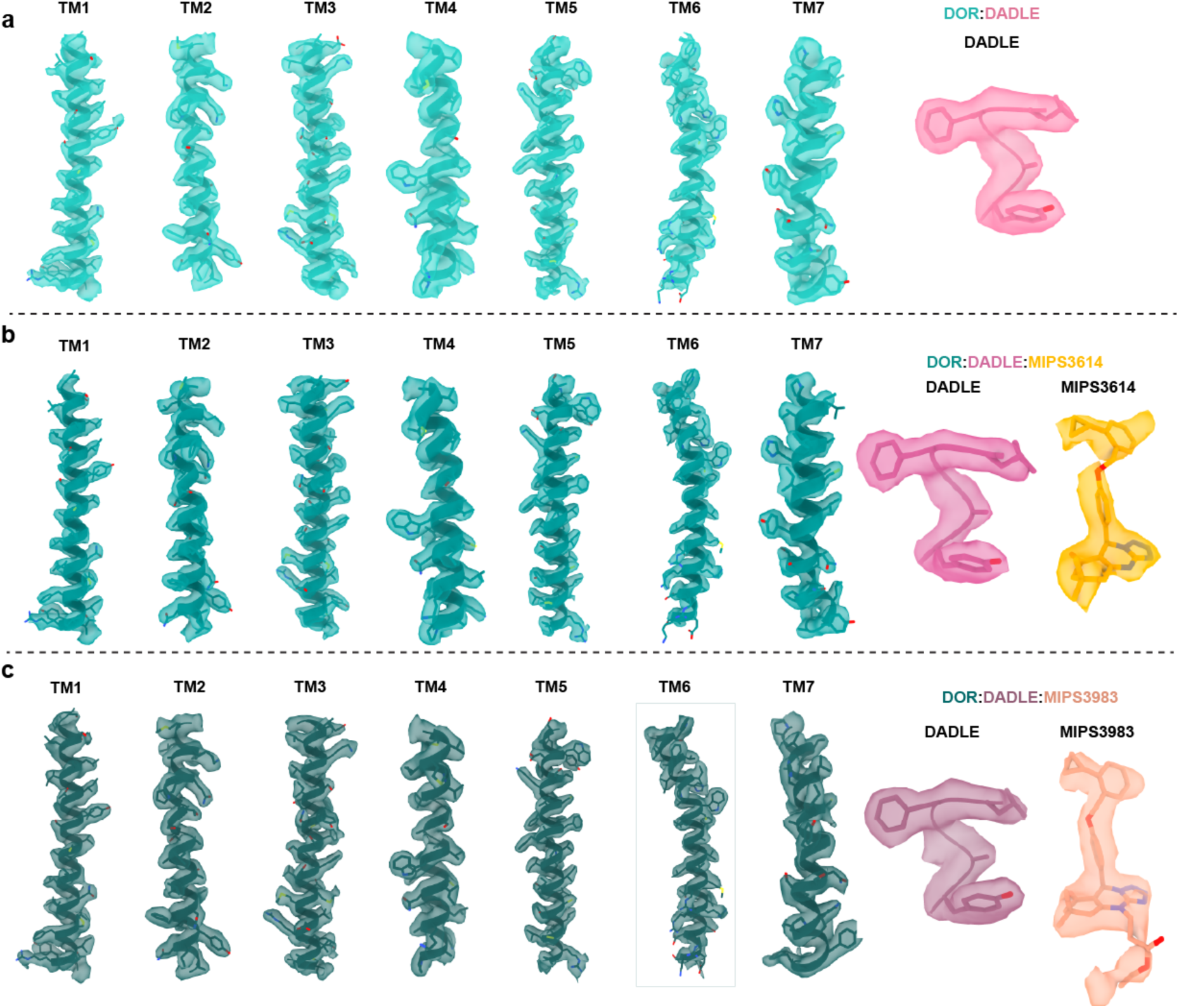
Comparison of cryo-EM density quality across δOR structures. (a) Cryo-EM density for the δOR-DADLE complex (2.0 Å resolution) showing individual transmembrane helices TM1-TM7 and the DADLE peptide. Shown is the receptor-focused map, contoured at 0.27. (b) Cryo-EM density for the δOR-DADLE-MIPS3614 complex (1.9 Å resolution) showing individual transmembrane helices TM1-TM7, the DADLE peptide and MIPS3614. Shown is the receptor-focused map, contoured at 0.42. (c) Cryo-EM density for the δOR-DADLE-MIPS3983 complex (1.9 Å resolution) showing individual transmembrane helices TM1-TM7, the DADLE peptide and MIPS3983. Shown is the receptor-focused map, contoured at 0.15.

**Supplementary Fig. 4.**
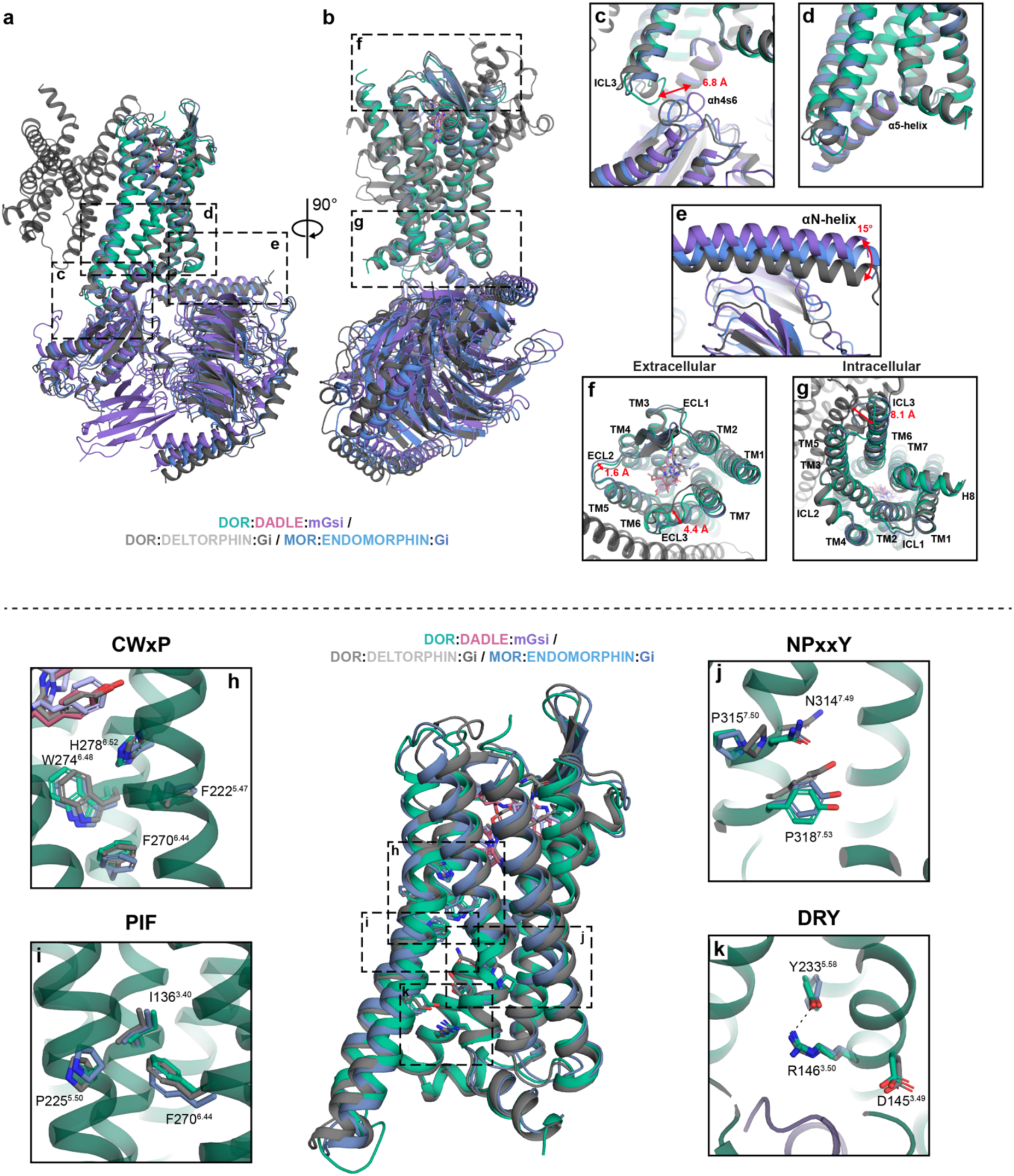
Structural alignments and activation motif analysis of δOR complexes. (a-b) Overall structural comparison of δOR-DADLE-mG_si_ (teal/blue) with δOR-Deltorphin-G_i_ (gray, PDB: 8F7S) and μOR-Endomorphin-G_i_ (light blue, PDB: 8F7R). (a) Side view showing the complete receptor-G protein complexes with excellent overall structural alignment. (b) Rotated view (90°) highlighting the conserved architecture across opioid receptor-peptide-G protein complexes. Dashed boxes (C, D, E, F, G) indicate regions shown in detailed panels. (c-g) Detailed views of structural differences between the receptor-G protein complexes. (c) Comparison of ICL3 and the Gα subunit. A large translation is observed for the αh4s6 loop (d) Comparison of the α5-helix interacting with the core of δOR. (e) Conformation of the N-terminal helix of mG_si_. (f) Extracellular view of the transmembrane domain showing differences in the position of ECL2 and ECL3. (g) Intracellular view highlighting the G protein coupling interface and differences in the position of ICL3 between structures. (h-k) Analysis of key GPCR activation motifs in their active conformations across the aligned structures. All structures show canonical active-state conformations. (h) The CWxP motif, (i) the PIF motif, (j) the NPxxY motif, and (k) the DRY motif.

**Supplementary Fig. 5.**
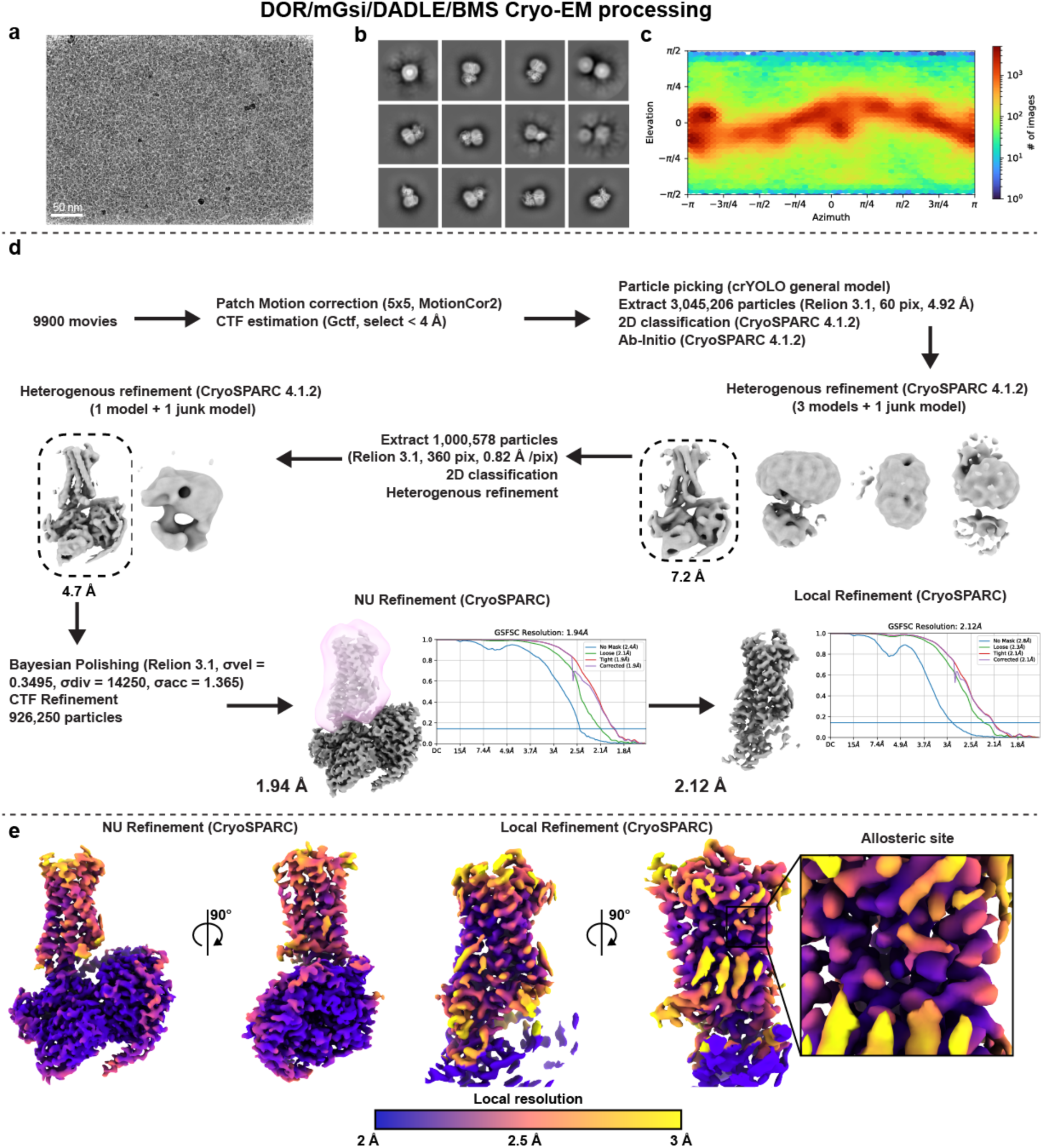
Cryo-EM data collection and processing workflow for δOR-mG_si_-DADLE-BMS-986187 complex. (a) Representative cryo-EM micrograph of the δOR-mG_si_-DADLE-BMS-986187 complex. Scale bar = 50 nm. (b) Gallery of 2D class averages from reference-free classification showing different orientations of the δOR-mG_si_-DADLE-BMS-986187 complex. (c) Angular distribution plot showing the coverage of particle orientations used in the final 3D reconstruction. (d) Comprehensive data processing workflow starting from 9,900 raw movies to a final 3D reconstruction with 926,250 particles. (e) Final 3D reconstruction colored by local resolution, showing two different views of the consensus (NU refinement) and receptor-focused (Local refinement) maps. The inset shows a detailed view of the allosteric binding site.

**Supplementary Fig. 6.**
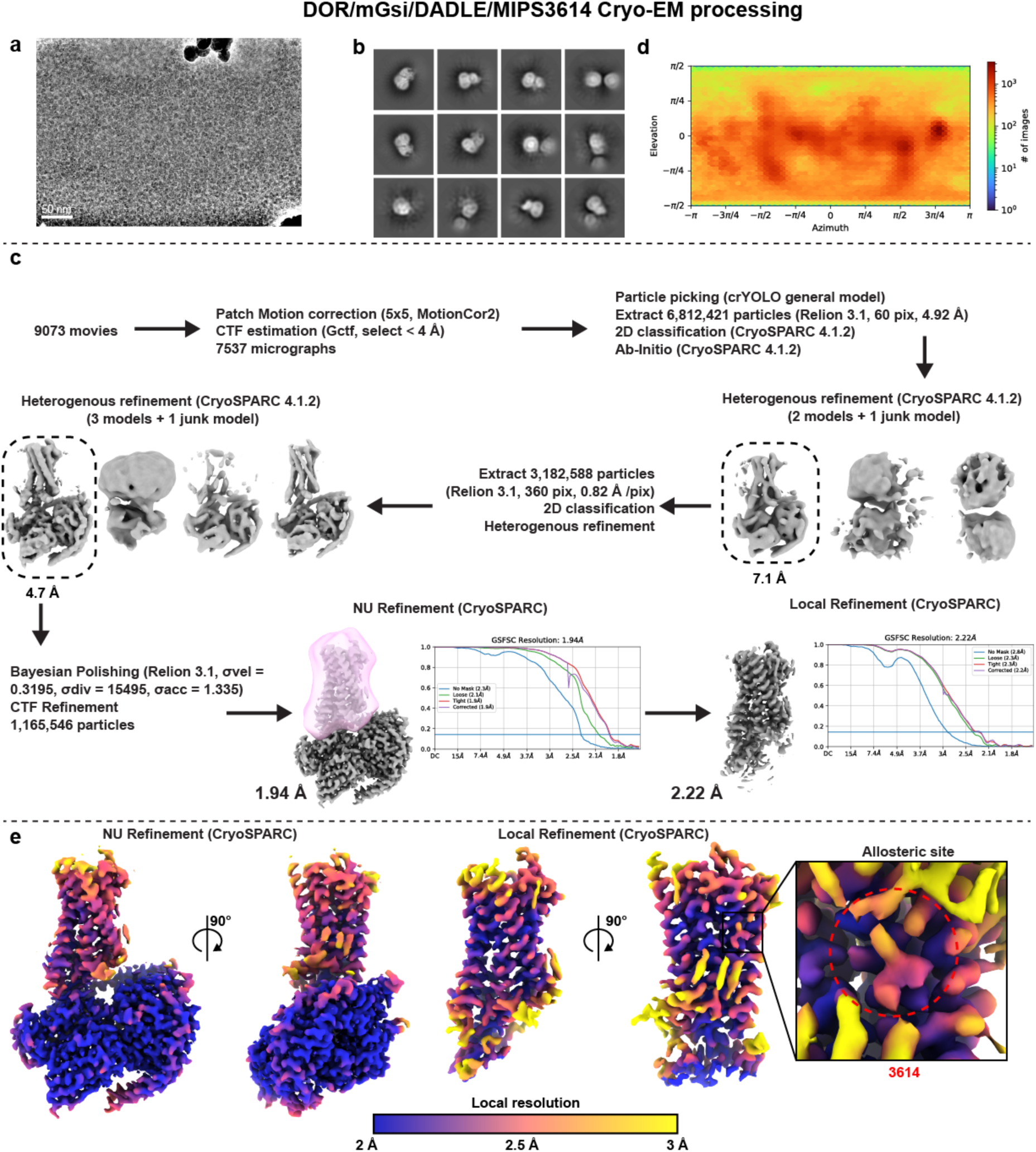
Cryo-EM data collection and processing workflow for δOR-mG_si_-DADLE-MIPS3614 complex. (a) Representative cryo-EM micrograph of the δOR-mG_si_-DADLE-MIPS3614 complex. Scale bar = 50 nm. (b) Gallery of 2D class averages from reference-free classification showing different orientations of the δOR-mG_si_-DADLE-MIPS3614 complex. (c) Angular distribution plot showing the coverage of particle orientations used in the final 3D reconstruction. (d) Comprehensive data processing workflow starting from 9,073 raw movies to a final 3D reconstruction with 1,165,546 particles. (e) Final 3D reconstruction colored by local resolution, showing two different views of the consensus (NU refinement) and receptor-focused (Local refinement) maps. The inset shows a detailed view of the allosteric binding site.

**Supplementary Fig. 7.**
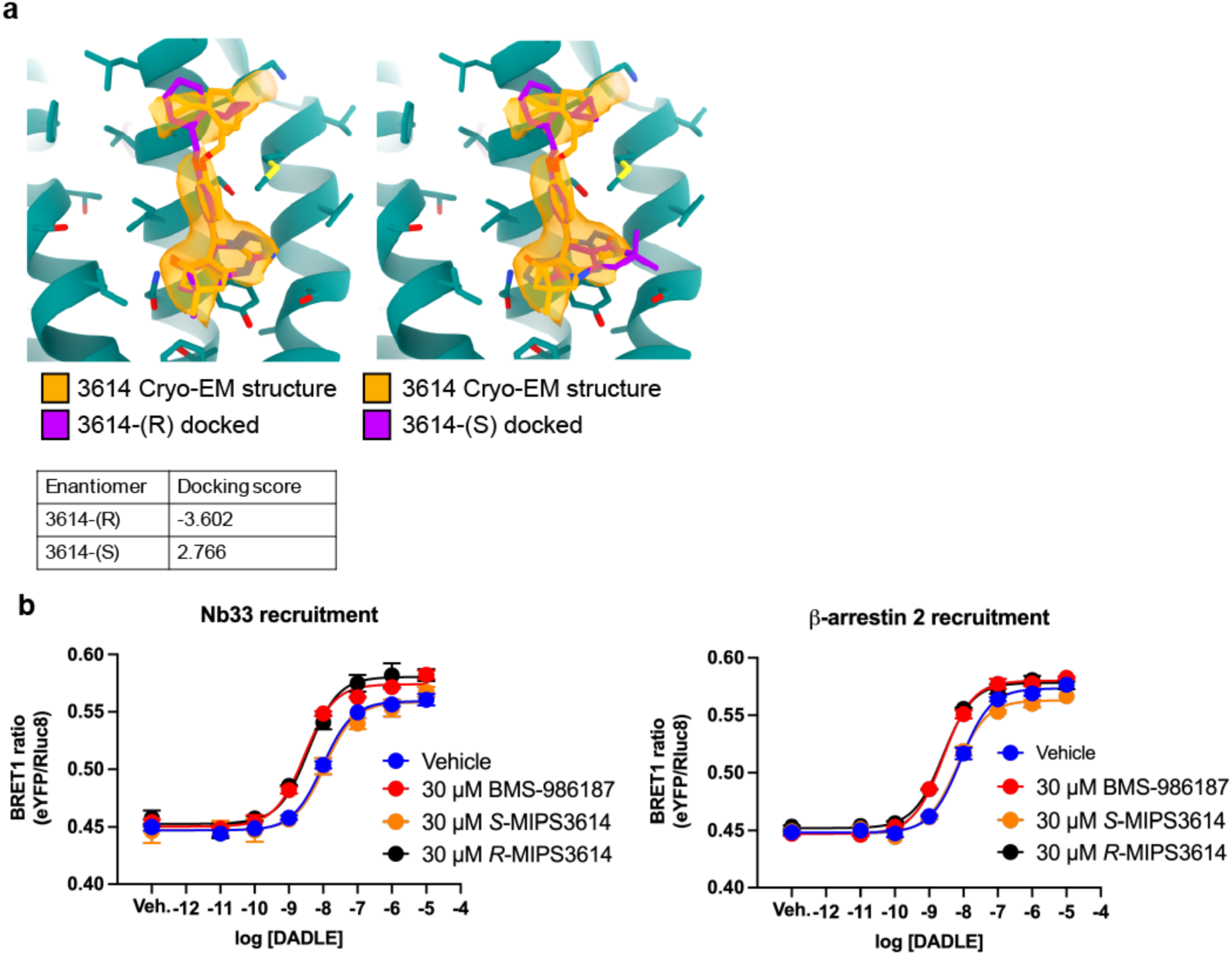
Structural and pharmacological evaluation of the *R*- and *S*-enantiomers of MIPS3614. (**a**) The binding site of MIPS3614 with the *R*-enantiomer docked on the left and the *S*-enantiomer docked on the right. The table below shows the docking scores. The fit to the cryo-EM map and docking scores are clearly better for the *R*-enantiomer. (**b**) Pharmacological characterisation of the *R*- and *S*-enantiomers of MIPS3614 compared to BMS-986187. Only one enantiomer showed activity in the Nb33 and β-arrestin 2 recruitment assay, which we assigned as the *R*-enantiomer, consistent with the cryo-EM structure. Data are representative of n=3 experiments, performed in duplicate.

**Supplementary Fig. 8.**
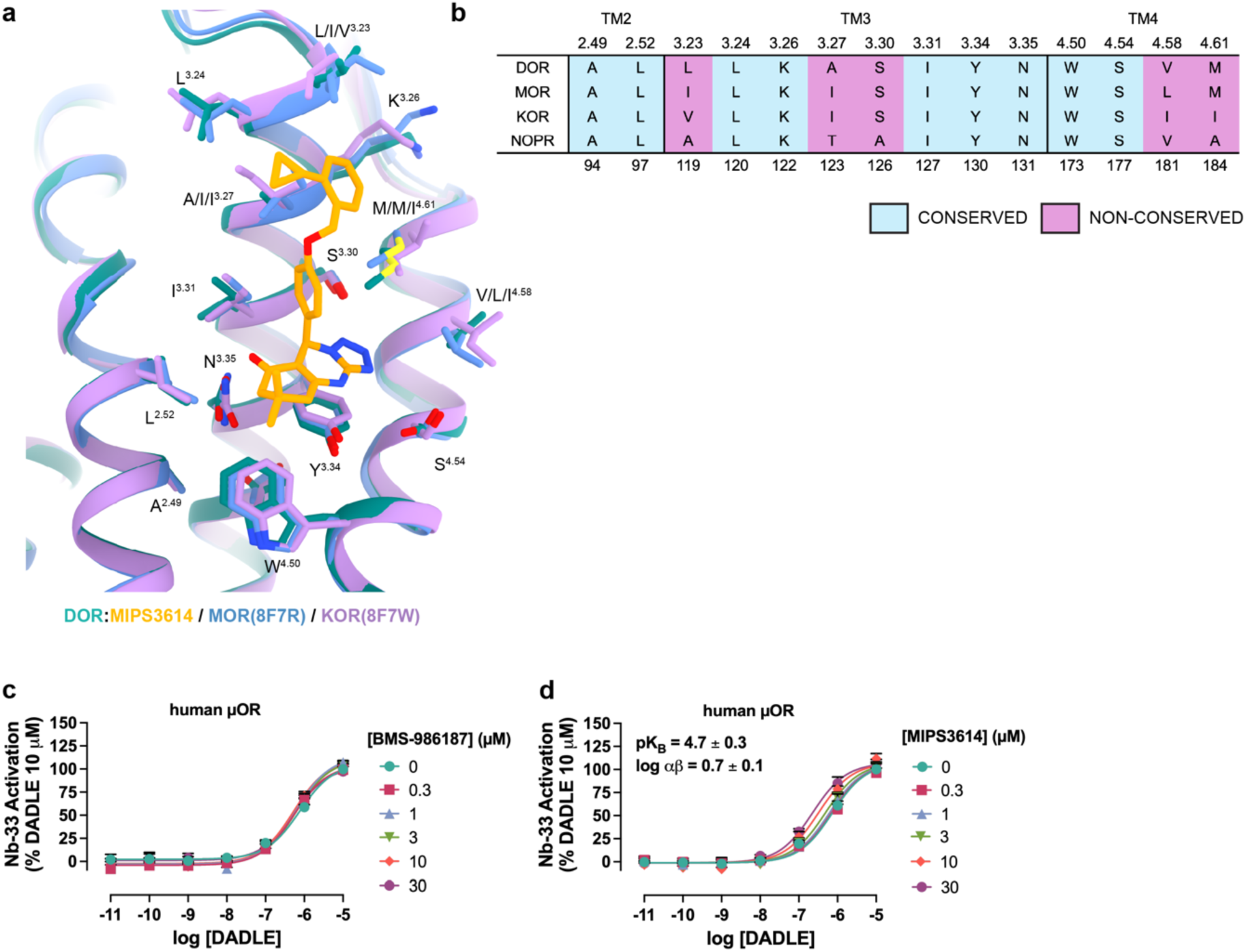
Conservation analysis of the MIPS3614 allosteric binding site across opioid receptor subtypes. (a) Structural overlay of the allosteric binding sites from δOR (teal, with MIPS 3614 in yellow), μOR (blue, PDB: 8F7R), and κOR (purple, PDB: 8F7W) showing the high degree of conservation in this region. Key residues involved in MIPS3614 binding are labelled with their Ballesteros-Weinstein numbers. The overlay demonstrates that the overall pocket architecture is similar across all three opioid receptor subtypes. (b) Sequence alignment table showing the conservation of allosteric binding site residues across the four opioid receptor subtypes (δOR, μOR, κOR, NOPR). Light blue boxes indicate fully conserved residues, while purple boxes highlight non-conserved positions that may contribute to subtype selectivity. (c) Pharmacology of BMS-986187 and MIPS3614 at the human µOR using a Nb33 recruitment assay (n=3). Not detectable allosteric activity of BMS-986187 was detected at µOR. In contrast, MIPS3614 had comparable activity at µOR and δOR (Fig. 2k).

**Supplementary Fig. 9.**
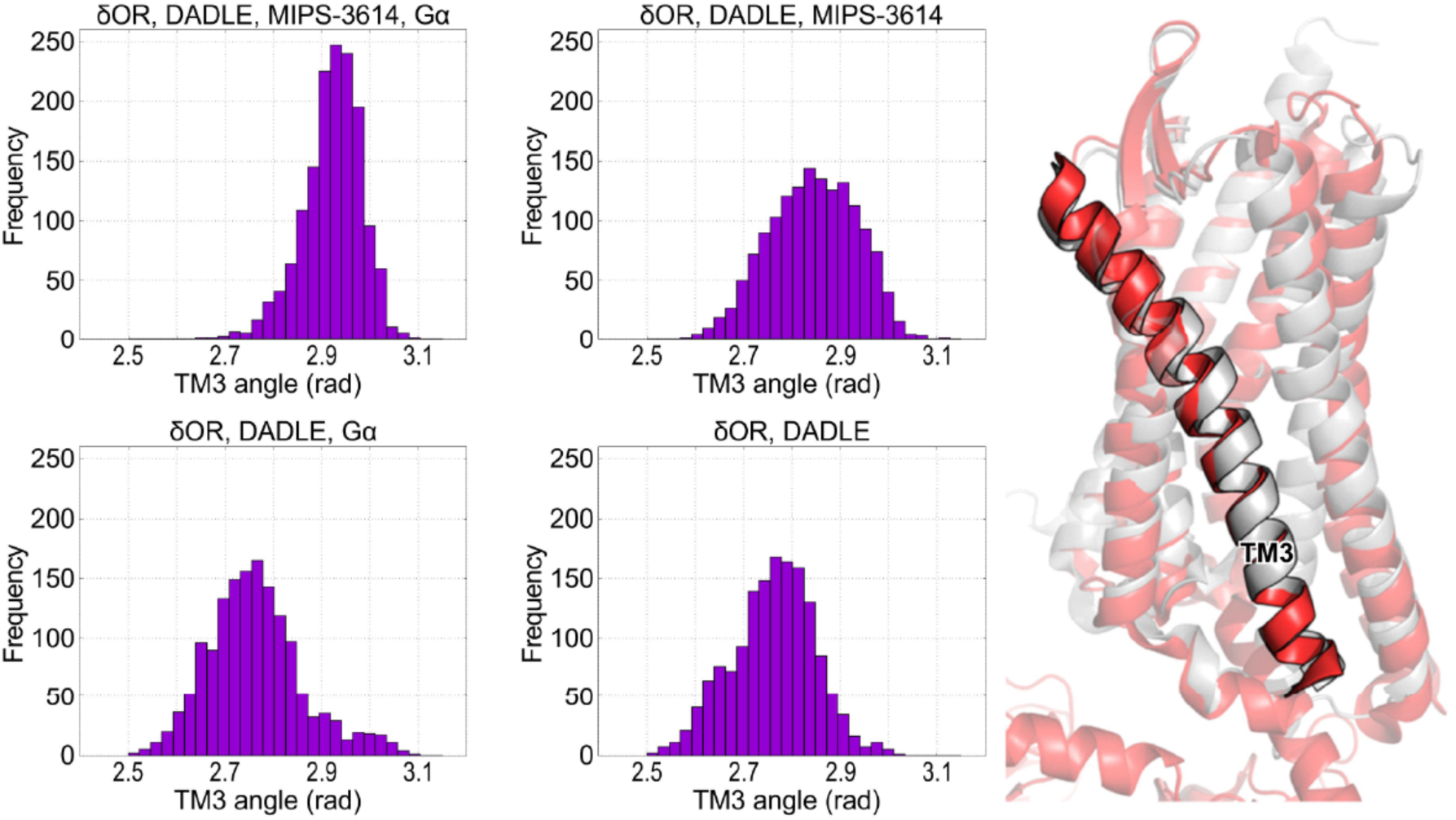
Molecular dynamics simulations reveal stabilization of TM3 bending angle by MIPS3614 binding. Histograms showing the distribution of TM3 bending angles (in radians) measured during 1 μs molecular dynamics simulations of different δOR complex states. The presence of MIPS3614, DADLE, or Gα protein individually or in combination stabilizes TM3 in a more bent conformation (higher angle values around 2.9 radians) compared to intermediate conformations observed with partial complexes. The structural representation (right) shows TM3 highlighted in black within the δOR structure (pink/gray), illustrating the helical bending measured in the simulations. This TM3 conformational change is associated with the active receptor state and correlates with the outward rotation of N131^3,35^ that enables MIPS3614 binding and sodium site disruption.

**Supplementary Fig. 10.**
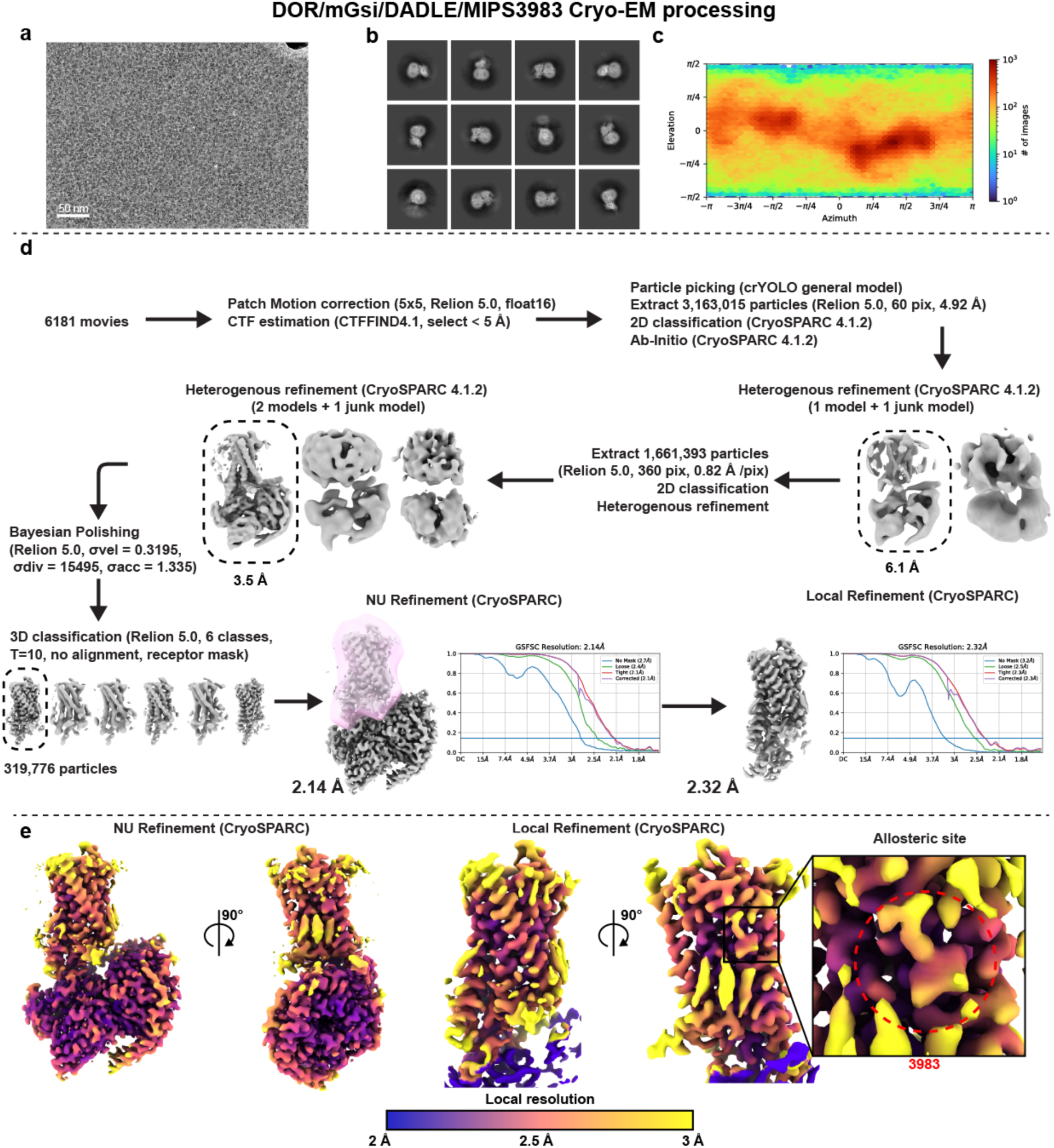
Cryo-EM data collection and processing workflow for δOR-mG_si_-DADLE-MIPS3983 complex. (a) Representative cryo-EM micrograph of the δOR-mG_si_-DADLE-BMS-MIPS3983 complex. Scale bar = 50 nm. (b) Gallery of 2D class averages from reference-free classification showing different orientations of the δOR-mG_si_-DADLE-MIPS3983 complex. (c) Angular distribution plot showing the coverage of particle orientations used in the final 3D reconstruction. (d) Comprehensive data processing workflow starting from 6,181 raw movies to a final 3D reconstruction with 319,776 particles. (e) Final 3D reconstruction colored by local resolution, showing two different views of the consensus (NU refinement) and receptor-focused (Local refinement) maps. The inset shows a detailed view of the allosteric binding site.

**Supplementary Fig. 11.**
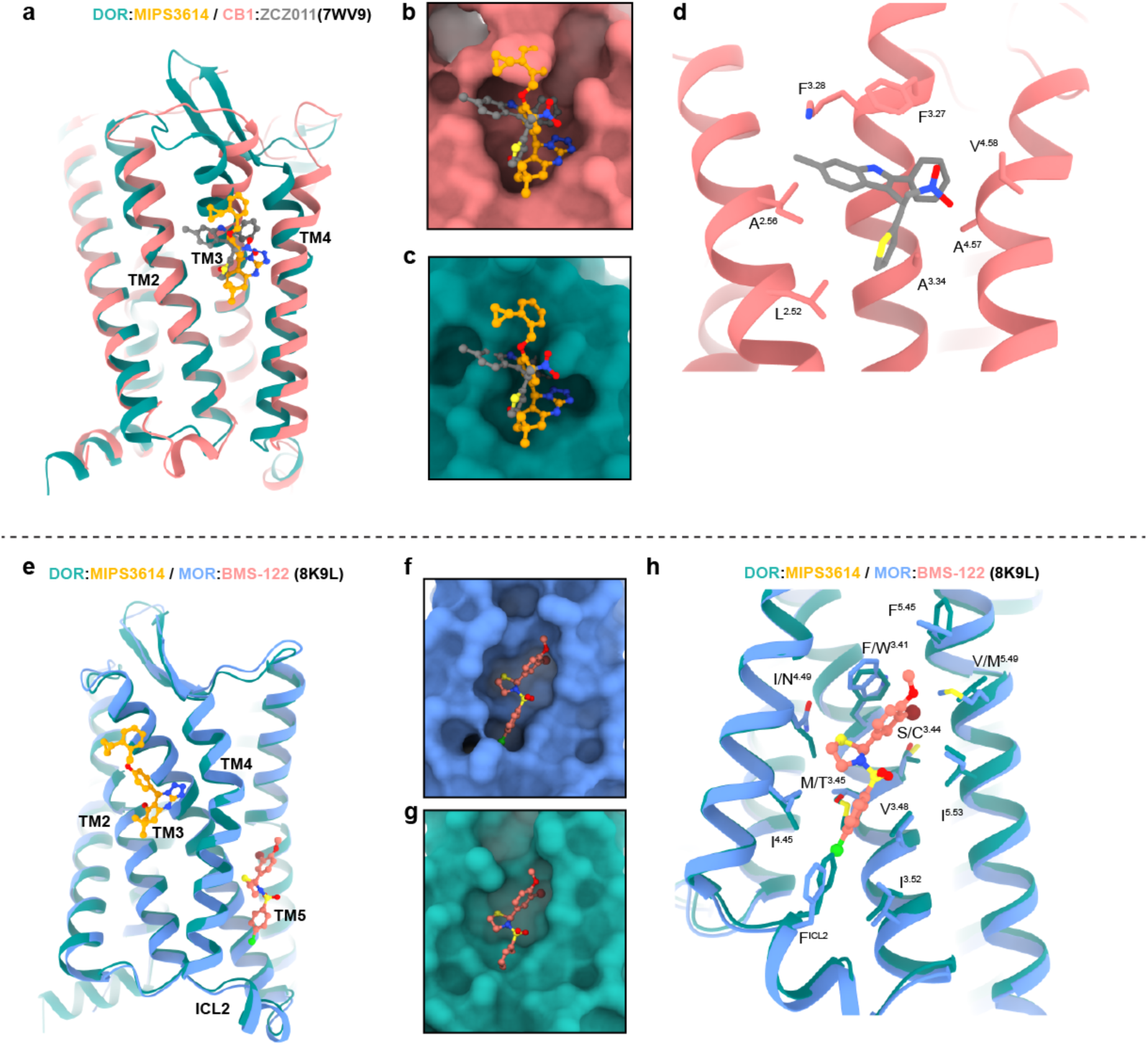
Comparison of the MIPS3614 allosteric binding site with the ZCZ011 binding site at the CB_1_ receptor. (a) Overview of the δOR-MIPS3614 structure (δOR in teal, MIPS3614 in orange) overlaid with the CB₁ receptor-ZCZ011 structure (CB₁ in pink, ZCZ011 in gray, PDB: 7WV9). The overlay shows the similar location of allosteric binding sites between TM2, TM3, and TM4 in both receptors, highlighting a potentially conserved allosteric site topology across different GPCR families. (b-c) The binding pockets of (b) CB1-ZCZ011 and (c) δOR-MIPS3614 are displayed as surfaces. The shape of the binding pockets differs significantly due to variations in hydrophobic interactions. (d) Detailed view of key residues forming the ZCZ011 binding site in CB₁. (e) Overview of the δOR-MIPS3614 structure overlaid with the μOR-BMS-122 structure (μOR in blue, PDB: 8K9L), showing the comparison between δOR and μOR allosteric binding sites. The allosteric binding sites between δOR and μOR are distinct. (f-g) Comparison of the binding surface of BMS-122 at (f) μOR and (g) δOR showing similar internal cavity architecture. (h) Detailed molecular view of the μOR-BMS-122 binding site showing key interacting residues. The binding pocket is relatively similar and conserved; however, μOR residue T^3.45^ is a methionine at δOR, which would likely clash with the binding of BMS-122.

**Supplementary Table 1.** Cryo-EM data collection, refinement, and validation statistics.

| | δOR/mG<sub>si</sub>/<br>DADLE | δOR/mG<sub>si</sub>/<br>DADLE/3614 | δOR/mG<sub>si</sub>/<br>DADLE/3983 | δOR/mG<sub>si</sub>/<br>DADLE/BMS<sup>#</sup> |
| --- | --- | --- | --- | --- |
| **Data Collection** |  |  |  |  |
| **EMD code\*** | **EMD-72825**<br>**(EMD-**<br>**72828)** | **EMD-72826**<br>**(EMD-**<br>**72829)** | **EMD-72827**<br>**(EMD-**<br>**72830)** | **EMD-72831**<br>**(EMD-72832)** |
| **PDB code** | **9YDP** | **9YDQ** | **9YDR** | **n/a** |
| **Micrographs** | **6008** | **9073** | **6181** | **9900** |
| **Electron Dose (e<sup>-</sup>**<br>**/Å<sup>2</sup>)** | **60** | **60** | **60** | **60** |
| **Voltage (kV)** | **300** | **300** | **300** | **300** |
| **Pixel size (Å)** | **0.82** | **0.82** | **0.82** | **0.82** |
| **Movie frames** | **60** | **60** | **60** | **60** |
| **Defocus range**<br>**(μm)** | **0.5-1.5** | **0.5-1.5** | **0.5-1.5** | **0.5-1.5** |
| **Refinement** |  |  |  |  |
| **Symmetry**<br>**imposed** | **C1** | **C1** | **C1** | **C1** |
| **Particles (final**<br>**map)** | **1,245,037** | **1,165,546** | **319,776** | **926,250** |
| **Resolution @0.143**<br>**FSC (Å)\*** | **1.95 (2.29)** | **1.94 (2.22)** | **2.14 (2.32)** | **1.94 (2.12)** |
| **CC<sub>map-model</sub>** | **0.90** | **0.88** | **0.89** |  |
| **Map sharpening B**<br>**factor (Å<sup>2</sup>)** | **-63** | **-53** | **-51** | **-51** |
| **Model Quality** |  |  |  |  |
| **R.M.S. deviations** |  |  |  |  |
| **Bond length (Å)** | **0.004** | **0.005** | **0.006** |  |
| **Bond angles (°)** | **0.713** | **0.819** | **0.836** |  |
| **Ramachandran** |  |  |  |  |
| **Favored (%)** | **98.64** | **98.55** | **99.23** |  |
| **Outliers (%)** | **0** | **0** | **0** |  |
| **Rotamer outliers**<br>**(%)** | **0.12** | **0.12** | **0.12** |  |
| **C-beta deviations**<br>**(%)** | **0.20** | **0.20** | **0.20** |  |
| **Clashscore** | **3.96** | **3.95** | **4.51** |  |
| **MolProbity score** | **1.18** | **1.18** | **1.23** |  |

**#BMS model not deposited to PDB** **\* Receptor focused refinement in parentheses**

**Supplementary Table 2.** Expression, Emax, and ΔpEC_50_ for δOR constructs from Figure 4.

| Construct | Expression (ELISA OD<sub>450nm</sub>) | | Emax | | $\Delta pEC_{50}$ | |
| --- | --- | --- | --- | --- | --- | --- |
| | mean ± SEM | p value | mean ± SEM | p value | mean ± SEM | p value |
| WT | 1.8 ± 0.2 | | 0.15 ± 0.02 | | 0.52 ± 0.04 | |
| A123<sup>3.27</sup>I | 0.5 ± 0.09 | <0.0001 | 0.12 ± 0.05 | 0.9993 | 0.24 ± 0.03 | 0.0158 |
| S126<sup>3.30</sup>A | 1.4 ± 0.3 | 0.4389 | 0.15 ± 0.04 | >0.9999 | 0.18 ± 0.01 | 0.0024 |
| S126<sup>3.30</sup>C | 1.6 ± 0.2 | 0.8508 | 0.11 ± 0.02 | 0.9358 | 0.30 ± 0.09 | 0.0774 |
| S126<sup>3.30</sup>F | 1.2 ± 0.2 | 0.0584 | 0.07 ± 0.01 | 0.1183 | -0.004 ± 0.13 | <0.0001 |
| Y130<sup>3.34</sup>A | 1.2 ± 0.1 | 0.0904 | 0.02 ± 0.01 | 0.0024 | ND | ND |
| N131<sup>3.35</sup>A | 1.1 ± 0.2 | 0.0302 | 0.021 ± 0.003 | 0.0021 | ND | ND |
| N131<sup>3.35</sup>S | 0.30 ± 0.04 | <0.0001 | 0.02 ± 0.01 | 0.0014 | ND | ND |
| N131<sup>3.35</sup>D | 1.6 ± 0.2 | 0.8815 | 0.06 ± 0.01 | 0.0908 | 0.02 ± 0.02 | <0.0001 |
| W173<sup>4.50</sup>F | 0.9 ± 0.1 | 0.002 | 0.057 ± 0.002 | 0.0555 | 0.24 ± 0.05 | 0.0165 |
| S177<sup>4.54</sup>A | 0.6 ± 0.1 | <0.0001 | 0.04 ± 0.01 | 0.0181 | ND | ND |
| S177<sup>4.54</sup>N | 0.6 ± 0.1 | <0.0001 | 0.015 ± 0.003 | 0.0012 | ND | ND |
| V181<sup>4.58</sup>F | 1.4 ± 0.1 | 0.4912 | 0.13 ± 0.06 | 0.9998 | 0.29 ± 0.15 | 0.1312 |
| M184<sup>4.61</sup>A | 1.7 ± 0.1 | 0.9998 | 0.18 ± 0.04 | 0.9816 | 0.22 ± 0.06 | 0.0074 |
| M184<sup>4.61</sup>F | 1.8 ± 0.1 | >0.9999 | 0.11 ± 0.02 | 0.8739 | 0.26 ± 0.02 | 0.0254 |
| Quad | 0.5 ± 0.1 | <0.0001 | 0.03 ± 0.01 | 0.0073 | ND | ND |

